# Sex-Specific Developmental Gene Expression Atlas Unveils Dimorphic Gene Networks in *C. elegans*

**DOI:** 10.1101/2023.08.05.552096

**Authors:** Rizwanul Haque, Sonu Peedikayil Kurien, Hagar Setty, Yehuda Salzberg, Gil Stelzer, Einav Litvak, Hila Gingold, Oded Rechavi, Meital Oren-Suissa

## Abstract

Sex-specific traits and behaviors emerge during development by the acquisition of unique properties in the nervous system of each sex. However, the genetic events responsible for introducing these sex-specific features remain poorly understood. In this study, we created a comprehensive gene expression atlas for both sexes of the nematode *Caenorhabditis elegans* across development. By comparing the transcriptome of pure populations of hermaphrodites and males from early larval stages to adulthood, we discovered numerous differentially expressed genes, including neuronal gene families like transcription factors, neuropeptides, and GPCRs. We identified INS-39, an insulin-like peptide, as a prominent male-biased gene expressed specifically in ciliated sensory neurons. We show that INS-39 serves as an early-stage male marker, facilitating the effective isolation of males in high-throughput experiments. Through complex and sex-specific regulation, *ins-39* plays pleiotropic sexually-dimorphic roles in temperature sensation, survival in cold temperatures, resilience against high hydrogen peroxide levels, and dauer entry, while also playing a shared, dimorphic role in early life stress. This study offers a comparative sexual and developmental gene expression database for *C. elegans*, which will facilitate research into the genetic regulation of the sexual development of other organisms. Furthermore, it highlights conserved candidate genes that may underlie the sexually-dimorphic manifestation of different human diseases.

## INTRODUCTION

Genetic sex introduces variation in phenotypic traits in sexually reproducing organisms. Sexual dimorphism, the phenotypic differences between the two sexes of a species, can be exhibited at various levels. These differences include variances in morphology^1^, sensory sensitivity^2^, social behavior^3, 4^, and disease progression or onset^5^. In essentially all sexually reproducing animals, from nematodes to humans, the nervous system undergoes sexual differentiation during a crucial developmental period, leading to changes in neuroanatomy, neural differentiation, synaptic connectivity, and physiology, which profoundly influence behavior^6–10^. This developmental shift is driven and accompanied by a sex-specific transcriptional program that has only been partially studied. Past work on developmental transcriptomes of various organisms, while capturing some universal features of sexual differentiation, were usually limited in scope, either focusing on one sex only^11, 12^ or ignoring sex altogether^13, 14^, covering limited developmental timepoints^15^, or focusing on non-neuronal tissues^11, 16, 17^.

To gain insight into the molecular and genetic mechanisms underlying the sexual component of nervous system development, we asked how sexual identity, neuronal identity, and developmental stage intersect to drive gene expression in a model organism. We addressed this question using the nematode *Caenorhabditis elegans*, due to the detailed anatomical and molecular understanding of the nervous system of both sexes and the extensive sexual dimorphism they exhibit^18, 19^ at the resolution of single identifiable neurons, connections, and behaviors^2, 20–22^. As most sexual differences arise late in development, *C. elegans* offers a unique opportunity to track how sex-specific characteristics emerge during neuronal development.

*C. elegans* is an androdiecious nematode, with hermaphrodites (XX) and males (XO). Males are rare (0.01%) in the standard lab strain, and no obvious morphological or molecular markers exist for large-scale male isolation before sexual maturation, a long-standing obstacle for systemic studies of sexual dimorphism or male development. Therefore, previous genome-wide expression studies in *C. elegans* have generally centered around just one sex, the hermaphrodite, and were limited to late larval stages or the use of pseudo males and relatively small sample sizes^23–28^. Traditional methods to purify adult males rely on manual picking^29^, mutations in *him* (*h*igh *i*ncidence of *m*ales)^30^ genes, or by size exclusion using mesh filters^31^, all of which are labor intensive and time-consuming. In a recent study^32^, large-scale preparation of L4 male larvae was achieved using auxin-induced degradation of a <u>d</u>osage <u>c</u>ompensation <u>c</u>omplex (DCC) component; nevertheless, male isolation at the early larval stages is still absent.

In the current work, we describe a gene expression atlas for the two sexes of *C. elegans* across development (https://www.weizmann.ac.il/sexpresso/). To achieve this, we developed a methodology that enables large-scale isolation of early larval males with high purity and then carried out whole animal RNA sequencing across all significant developmental stages for both sexes. Our findings reveal a multitude of sexually-dimorphic differentially expressed genes, including neuronal gene families such as transcription factors (including DM domain and homeobox genes), neuropeptides, and GPCRs. We validated our findings using multiple approaches and identified an early-stage marker for males. Comprehensive anatomical localization and functional studies of one of the top hit genes, the insulin-like neuropeptide INS-39, revealed sexually-dimorphic expression and functions in the two sexes. We also discovered a complex and sexually-dimorphic regulation system for this gene in the two sexes. Together, the extensive database of dimorphic gene expression across development provided by this study will serve as a source for determining the functions of specific genes in both sexes. Finally, our study also draws attention to conserved candidate genes that might be linked to the sexually dimorphic manifestations of human diseases.

## RESULTS

### Large scale-male isolation in early larval stage of *C. elegans*

To obtain a male-enriched worm population we exploited a temperature-sensitive mutation in a dosage compensation complex gene detrimental only to hermaphrodites. The dosage compensation complex (DCC), a specialized regulatory mechanism that downregulates gene expression from the two X chromosomes, is only formed in *C. elegans* hermaphrodites (males carry just one X chromosome) (Figure 1A)^33^. Consequently, defective DCC will result in a lethal overdose of gene expression from the X chromosomes only in hermaphrodites^34–36^. We used a temperature-sensitive allele (*y1*) in the DCC gene *dpy-28*^37^ with a *him-8* mutation that spontaneously leads to a high incidence of males (Figure 1B). As expected, at the restrictive temperature (25°C), most XX hermaphrodites died as embryos or L1 larvae, whereas the XO males were unaffected. However, even at the restrictive temperature, 20% of XX hermaphrodites were still viable (Figure 1B). We observed that the L1 XX hermaphrodites that did hatch at the restrictive temperature were severely impaired, displaying body defects and defective locomotion. We utilized this sex-specific phenotype and improved male purity by slicing out and removing the immobile drop of hermaphrodites from the plate (Figure S1, see methods). Using this modified protocol, we improved male enrichment to over 98% (Figure 1B).

**Figure 1:**
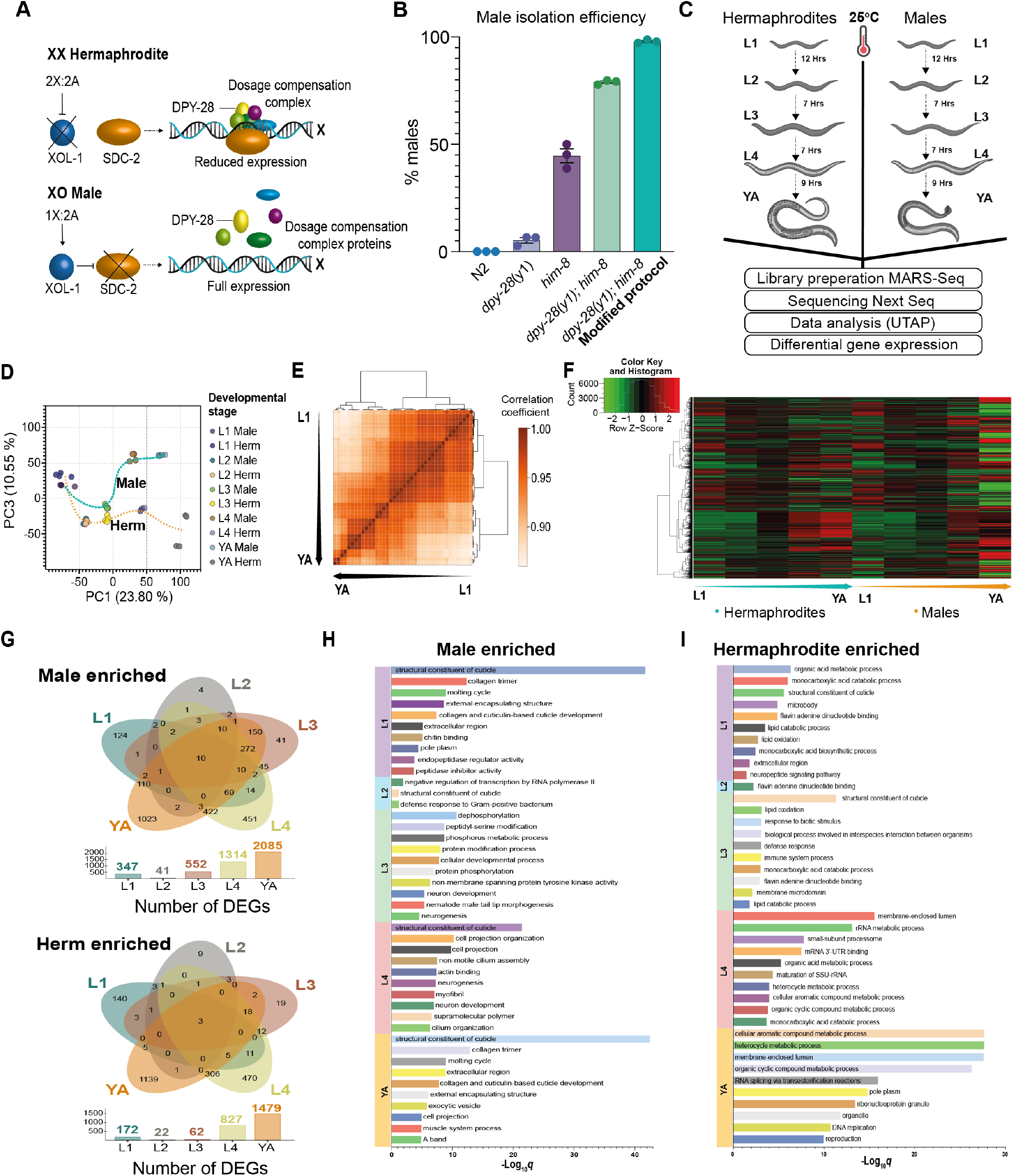
Large-scale isolation of pure male populations enables profiling of whole animal transcriptomes across all developmental stages. (A) A model for *C. elegans* dosage compensation complex (DCC) forming on the X chromosome only in hermaphrodites (image modified from^51^). (B) Temperature-sensitive mutation (*y1*) in the dosage compensation gene, *dpy-28*, facilitates large-scale male isolation. Quantification of male percent at YA developmental stage in N2, *dpy- 28*(*y1*), *him-8*(*e1489*), *dpy-28*(*y1*);*him-8*(*e1489*) and *dpy-28*(*y1*);*him-8*(*e1489*) (modified protocol) at 25°C with 3 biological replicates, n=100 worms per group. Error bars are standard error of the mean (SEM). (C) Schematic of experimental design and RNA-seq procedure (see methods). (D) Principal component analysis projection of expression patterns for the five developmental stages of the two sexes. Each circle represents individual samples. The progression of developmental stages is shown as a dotted line for each sex (cyan, males, and orange, hermaphrodites). The % variance, out of the total original variance in the high-dimensional space, spanned by the first and third PCs is indicated on the x and y axis, respectively. (E) Sample correlations. Hierarchical clustering of all sample types based on their RNA expression profiles. Off-diagonal entries denote the Pearson correlation between the expression profiles of two different sample types. Here, the sample classes cluster well, showing the highest intra-group correlations (developmental stage and sex). (F) Hierarchical clustering of differentially expressed genes (by DESeq2) using the genes expression values (rlog transformed counts (rld)) on a per-developmental stage basis, using the thresholds for significant differential expression as padj ≤ 0.05, |log2 fold change| >= 1 and basemean >= 5. Each row represents a gene. The arrow indicates the progress of developmental stages and results are clustered only by rows. (G) Venn diagrams showing genes enriched in males (top) and hermaphrodites (bottom) across all developmental stages. (H-I) Top 10 GO term enrichments for males enriched (H) and hermaphrodites enriched (I) set of genes for each developmental stage. The significance of the enrichment at a particular stage was determined using *q*-value threshold of 0.1.

To determine the suitability of the isolated *dpy-28* males to serve as a source for wild-type male RNA sequencing, we assayed their morphology, locomotion and mating behavior, and compared them to wild-type males. *dpy-28* males were indistinguishable from *him-8* control males in multiple parameters and behaviors, including body size, locomotion, and mating efficiency (Figure S2). Relatedly, previous studies have shown that RNAi against *dpy-28* exclusively affects hermaphrodites, and not males, in lifespan and dauer arrest^38, 39^. Therefore, we concluded that, unlike *dpy-28* hermaphrodites, *dpy-28* males develop normally into healthy adults, and can thus serve as an opposite-sex counterpart to wild-type hermaphrodites for comparative transcriptome analyses.

### Genetic sex and development converge to generate unique whole animal transcriptomes

Having established a reliable procedure to obtain pure male populations starting from early juvenile stages, we sought to carry out whole-animal RNA-seq transcriptomics in both sexes and throughout all developmental stages. To achieve this, we used a modified bulk version of the single-cell MARS-seq protocol. This modified approach is a high-throughput, low-input 3’- mRNA-seq method, which enhances the quality of library preparation for more accurate gene expression profiling^40, 41^. We produced RNA-seq profiles from five distinct developmental stages (L1, L2, L3, L4, YA) for both sexes, totaling 40 samples (10 groups with four biological repeats) (Figure 1C). Male percentages, raw read counts, and RINe scores per sample are presented in Supplementary Table S1. PCA analysis and hierarchical clustering of pairwise sample Pearson correlations grouped RNA profiles from biological repeats with high confidence, exhibiting the highest correlations for intra-group samples (Figure 1D-E). Notably, when our samples were separated by stage on principal component 1 (PC1), and by sex on principal component 3 (PC3), samples from both sexes clustered together before the onset of sexual maturation (i.e., L1-L3 stages), and robustly diverged after sexual maturation (L4-YA stages), reflecting the dramatic anatomical and physiological changes that occur during sexual maturation. We detected expression data for 14185 genes (Figure 1F), among which 4698 genes (33%) showed dimorphic expression in either developmental stage, including 2474 known genes and 2224 predicted novel genes. The expression data revealed 519 (L1), 63 (L2), 614 (L3), 2141 (L4), and 3564 (YA) genes that were differentially or dimorphically expressed (Figure 1G, for raw expression data see Supplementary Table S2). Overall, there were more male-biased differentially expressed genes (DEGs) (henceforth called male enriched genes (MEGs)) than hermaphrodite-biased DEGs (hermaphrodite enriched genes (HEGs)) at all developmental stages, suggesting an inherent sex-dependent bias of gene expression (Figure S3A).

To validate our datasets, we confirmed the expected expression pattern of known markers necessary for the development of sex-specific features, which typically appear during the third larval stage^42^. *pkd-2*, for instance, is expressed in the cilia of three different types of male sensory neurons^43^*, K09C8.2* in the seminal vesicle and vas deferens^44^, *clec-207* in the vas deferens^27^, and *Y39B6A.9* in male spicule muscle^28^, *R01E6.5* in male spicule neuron^28^ and *ram-5* in the ray neurons^45, 46^. For hermaphrodites, we used *meg-1,* which is expressed in the proximal germline^47^, and *vit-2* in the vesicles of the oocyte^48^. Our data coincides with these genes’ reported stage-specific expression pattern, which shows essentially no to minimal expression in males/hermaphrodite before sexual maturation and increases drastically during L4 when sex-specific structures emerge (Figure S3C-J).

Next, we analyzed our datasets using jvenn^49^ and discovered that only 10 Male-enriched genes and 3 hermaphrodite-enriched genes remained enriched sex-specifically throughout all developmental stages (Figure 1G, Supplementary Table S3). To gain further insight, WormBase gene ontology (GO) enrichment analyses were applied to the DEGs for each developmental stage and sex^50^. Male-enriched genes at early development were significantly associated with cuticle development and at later stages with neurogenesis and molting (Figure 1H). In contrast, hermaphrodite-enriched genes at early development were significantly associated with metabolic pathways and at later stages with reproduction and gamete formation (Figure 1I). Taken together, profiling of whole animal transcriptomes across all developmental stages reveals a distinct sex-specific developmental plan, before any morphological sexual features arise.

### Distinct neuronal gene families are regulated in a sexually dimorphic manner across development

From worms to humans, the nervous system undergoes extensive transcriptional and functional transformation upon sexual maturation^19, 22, 52^. We thus searched specifically for neuronal gene families that display sexually dimorphic expression in our dataset. We examined transcription factor families^53^, including DM domain genes^54^ and homeobox genes. 218 TFs were differentially expressed in at least one developmental stage (Supplementary Table S4, Top 25 represented in Figure 2A), including previously known male-specific TFs^46, 55^ such as *mab-3*, *mab-5*, *egl-5*, *dmd- 3* and the hermaphrodite-specific TFs^56, 57^ *lin-13* and *lin-15.* This list uncovered a pool of TFs with temporal and sexual specificity during development. While only a few TFs were differentially expressed at early development (Figure S3B), we found over 100 that were differentially expressed in young adults (Supplementary Table S4). 73 TFs were specifically enriched in males, and 74 were specifically enriched in hermaphrodites. Intriguingly, we found just two TFs in our data, *ces- 2* and *mab-3,* that were consistently higher in males across all developmental stages (Figure 2A- D, Figure S4E). The DM (**D**oublesex/**M**AB-3) domain genes regulate sexual development across evolution and are integral players in sexual development and its evolution in many metazoans^58^. We found three DM domain genes to be expressed dimorphically, namely *mab-3*, *dmd-4,* and *dmd- 3* (Figure 2D and Supplementary Table S4), all of which have been previously shown to play sex- specific roles^46, 59, 60^. We also noticed that most DMDs are regulated temporally in both sexes, suggesting they might play stage-specific roles in shared developmental processes.

**Figure 2:**
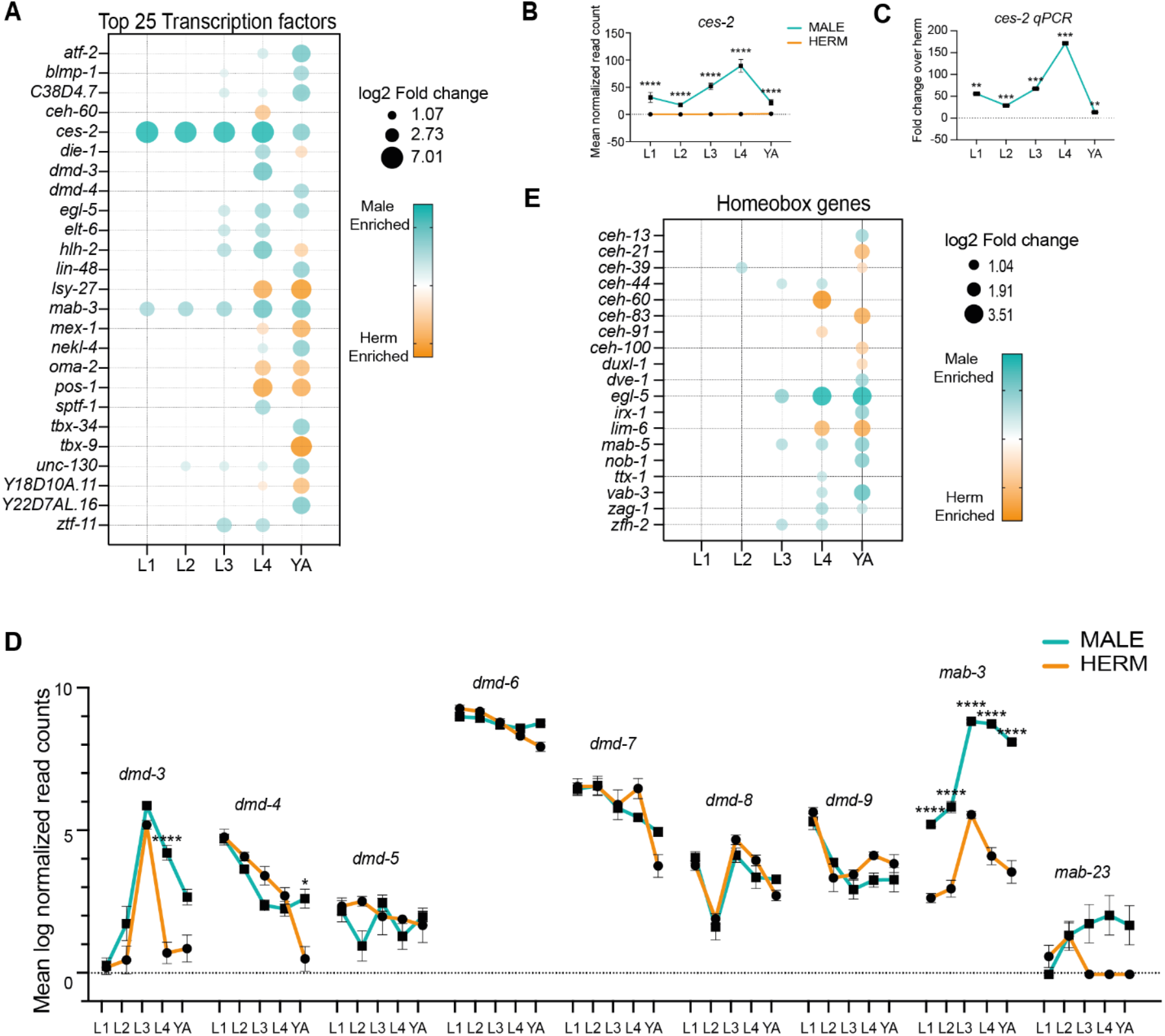
Selected gene families with sexually dimorphic regulation. (A) Bubble plot representation of top 25 transcriptional factors out of 218 differentially expressed in any of the five developmental stages of the two sexes. Bubble size represents log2 of fold change in expression of that gene (only genes that passed the filter padj <= 0.05, |log2 fold change| >= 1 and basemean >= 5 are plotted), bubble color (cyan: males and orange: hermaphrodites) represents enrichment in either sex. TFs were mined using wTF2.2, an updated compendium of *C. elegans* transcription factors^53^. (B) Normalized RNA-seq expression values of *ces-2* across all developmental stages. (C) Real-time qPCR analysis of *ces-2* mRNA (*ces-2* mRNA male expression normalized to hermaphrodite expression) across all developmental stages. (D) Log2 normalized expression values of differentially expressed DM domain genes across development. (E) Bubble plot representation of homeobox genes differentially expressed in any of the five developmental stages of the two sexes. Bubble size represents log2 of fold change in expression of that gene (only genes that passed the filter padj <= 0.05, |log2 fold change| >= 1 and basemean >= 5 are plotted), bubble color (cyan: males and orange: hermaphrodites) represents enrichment in either sex. Error bars are standard error of the mean (SEM). In (C) p-values were calculated by t-test for each comparison performed. In (B, D) adjusted p-values were calculated by Wald test for each comparison performed by DESeq2^77^, **** p < 0.0001, *** p < 0.001, ** p < 0.01, * p < 0.05, ns-non-significant.

Homeobox genes regulate various aspects of development and specification of neuronal identity in *C. elegans* and across evolution^61^. It was recently shown that neuronal diversity in *C. elegans* can be fully described by unique combinations of the expression of homeobox genes^62–64^. Thus, dimorphism in homeobox gene expression might imply that there is sexual dimorphism also in the code that defines the neuronal identity features. We found that 19 of the homeobox-containing genes were sexually dimorphic across various stages of development (Figure 2E, Supplementary Table S4). Our data corroborate the high male expression of *egl-5, vab-3*, *mab-5, ceh-13* and *nob- 1*, required for several aspects of male sexual differentiation, like the formation of male-specific sensory organs, sex muscle differentiation, or gonadal development^65–69^. Additionally, we found that *ttx-1, zfh-2, zag-1,* previously shown to be involved in neurogenesis^70–72^ and *irx-1* in synapse elimination^73^, are expressed significantly higher in males during sexual maturation (Figure 2E) and can thus be potentially involved in male-specific neurogenesis/rewiring which have not been explored before. Similarly, we found extensive sexual dimorphism throughout development for neuronal terminal differentiation genes, such as K channels^74^, ligand-gated ion channels^74^, ionotropic receptors^74^, synaptic vesicle genes^75^, and nuclear hormone receptors^76^ (Figure S5A-E, Supplementary Table S4). In summary, our data provide a rich resource for the factors that may drive male neuronal identity and its functional landscape.

### Analysis of differentially-regulated neuropeptides reveals the insulin-like peptide INS-39 to be highly sexually dimorphic

Neuropeptides constitute a vast class of signaling molecules in the nervous system of many groups of animals, yet despite their prevalence, dimorphic functions have been assigned only to few neuropeptides^78–84^. We thus focused on the large, extended family of neuropeptides in *C. elegans* and their cognate GPCR receptors. Among the receptors, *srj-49* caught our attention for being extremely dimorphic as it is upregulated in males starting already at the L1 stage and throughout all subsequent developmental stages (Figure 3A). Real-time qPCR analysis validated that *srj-49* mRNA is absent in hermaphrodites at all stages, while it is highly expressed in males (Figure S4A- B). However, we observed no detectable SRJ-49 protein neither by using an *srj-49*p::GFP multi- copy fosmid reporter nor by a single-copy *srj-49*::SL2::GFP::H2B CRISPR reporter (Figure S4C- D). We propose that *srj-49* may be a non-coding RNA that has male-specific functions.

**Figure 3:**
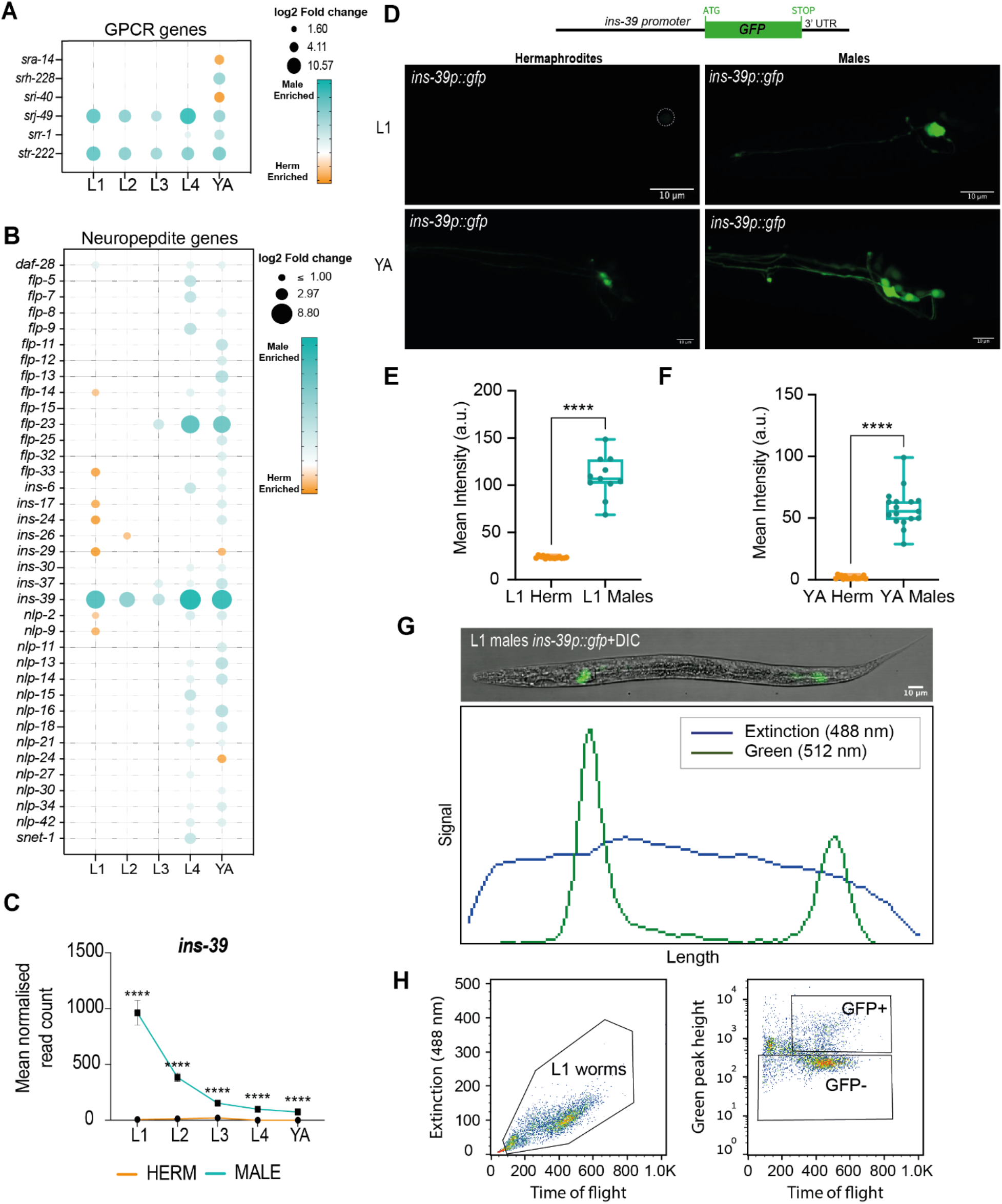
Neuropeptide superfamily genes with sexually dimorphic regulation. (A-B) Bubble plot representation of GPCRs (A) and neuropeptide genes (B) differentially expressed in any of the five developmental stages of the two sexes. Bubble size represents log2 of fold change in expression of that gene (only genes that passed the filter padj <= 0.05, |log2 fold change| >= 1 and basemean >= 5 are plotted), and bubble color represents enrichment in either sex (male enrichment in cyan, hermaphrodite enrichment in orange). (C) Normalized expression values of *ins-39* across all developmental stages in both sexes (cyan: males, orange: hermaphrodites). Error bars are standard error of the mean (SEM). (D) Top, schematic of the *ins-39p::gfp* expression reporter used in this study. Bottom, representative confocal micrographs of the expression pattern of the *ins-39p::gfp* reporter in head neurons at L1 and YA stage in both sexes (bottom). Scale bars represent 10µm. (E-F) Quantification of *ins-39p::gfp* fluorescence intensity from D in head neurons in both sexes at L1 (E) and YA stage (F). a.u, arbitrary units. Vertical bars in the box-and-whiskers graph represent the median, with dots showing all points from min to max. n = 11-14 animals for (E) and n = 17 animals for (F) per group. (G) *ins-39p::gfp* enables male isolation using flow cytometry. Top, representative confocal micrographs of a whole L1 male expressing *ins-39p::gfp*. Bottom, extinction and fluorescence profiles from *ins-39p::gfp* L1 males on the BioSorter profiler. The worm is oriented with its head towards the left side of the frame. The BioSorter profiler accurately captures the GFP intensity profile from the *ins-39p::gfp* expressing neurons (green channel), providing an effective means to sort L1 males from L1 hermaphrodites. Scale bars represent 10µm. (H) Gating profiles for *ins-39p:gfp* expressing worms based on time of flight in the flow cytometer flow cell and extinction at 488 nm. Gate region selected for L1 worms (time of flight vs. extension, left panel), and gate region selected for GFP- and GFP+ (time of flight vs. GFP intensity peak height, right panel). We performed a Mann-Whitney test in (E, F). In (C), adjusted p-values were calculated by Wald test for each comparison performed by DESeq2^77^, **** p < 0.0001.

The *C. elegans* genome encodes 154 known neuropeptide genes, 40 genes belong to the insulin- like family of peptides, 31 genes are FMRFamide-related peptides, and 83 genes encode non- insulin, non-FMRFamide-related neuropeptides^85^. Our analysis shows that 37 neuropeptide genes exhibit dimorphic expression with a strong bias toward male enrichment (Figure 3B). We found that only 12 neuropeptide genes exhibited dimorphic expression before the L4 stage, indicating that most neuropeptide-dependent signaling pathways diverge only after sexual differentiation. Interestingly, we found a single neuropeptide gene, *ins-39* insulin/IGF1 hormone, that was consistently expressed higher in males during all developmental stages (Figure 3B-C). We validated the higher expression of *ins-39* in males using a transcriptional *ins-39p::gfp* reporter array. Remarkably, adult males exhibited 32-fold higher mean fluorescence intensity than hermaphrodites (Figure 3D-F), and L1 hermaphrodites showed essentially no fluorescence at all (Figure 3D, E). The male-specific expression of *ins-39p::gfp* enabled us to distinguish L1 males from hermaphrodites both manually or automatically using a complex object sorter (COPAS)^86^ (Figure 3G-H), thus providing a needed early-stage male-specific marker as a powerful tool for high throughput isolation of males.

### INS-39 is highly enriched in a subset of sex-shared sensory neurons in males

INS-39 has recently been reported to be expressed in AFD neurons^87^, but its male pattern suggests a broader expression. To determine the complete expression pattern for INS-39, we generated a nuclear-localized CRISPR/Cas9-engineered reporter allele [*ins-39*::SL2::GFP::H2B] (Figure 4A). This reporter revealed a striking dimorphism across all stages: while hermaphrodite expression is low and restricted, males exhibit higher and broader neuronal GFP expression throughout development (Figure 4B-C). We used “NeuroPAL” (<u>Neuro</u>nal <u>P</u>olychromatic <u>A</u>tlas of <u>L</u>andmarks) to identify cell types using color barcodes^88, 89^. We found that in young adult hermaphrodites, INS- 39 was specifically expressed in two sets of neurons, AFD and ASK. In contrast, in males, INS-

**Figure 4:**
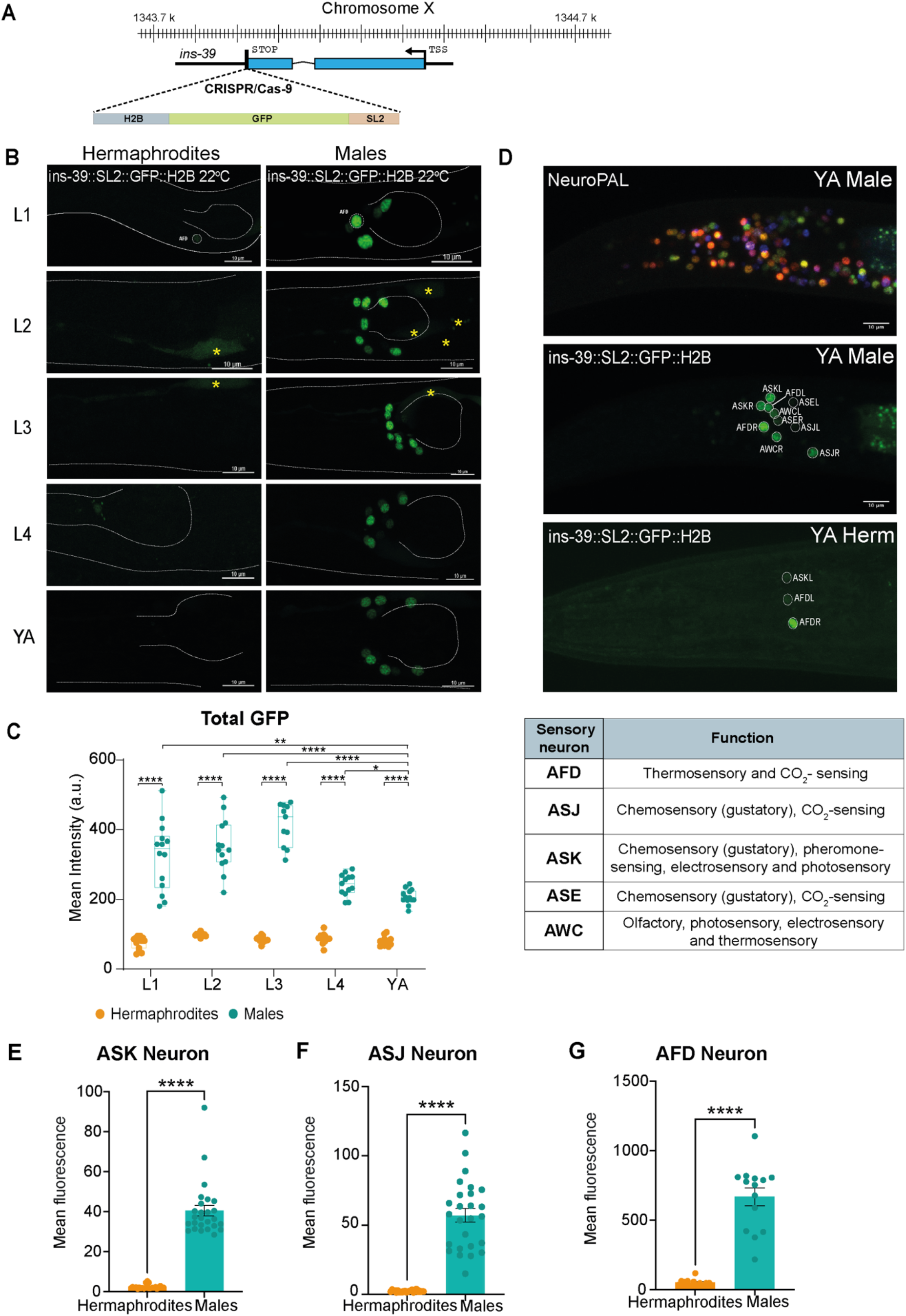
Sexually-dimorphic expression pattern of INS-39. (A) Schematic of the CRISPR/Cas9-edited *ins-39*(*syb4915*[*ins-39*::SL2::GFP::H2B]) reporter. (B) Representative confocal micrographs of the expression pattern of *ins-39*(*syb4915)* reporter in *C. elegans* across all developmental stages in both sexes. The head and pharynx are outlined in a white dashed line. Co-injection marker is indicated by yellow asterisk mark. Scale bars represent 10µm. (C) Quantification and comparison of mean total head GFP intensity of the *ins-39*(*syb4915)* reporter expression. n =11-14 worms per group. Vertical bars in box-and-whiskers graph represent the median, with dots showing all points from min to max (with dots depicting individual data points). (D) Representative confocal micrographs showing genetically encoded multi-colored neuronal nuclei of a NeuroPAL worm (top) used to identify the neurons expressing the *ins-39*(*syb4915)* reporter in young adult males (middle) and hermaphrodites (bottom). The table below lists the identified sensory neurons’ established functions (www.wormatlas.org). Scale bars represent 10µm. (E-G) Quantification of mean GFP intensity in ASK (E), ASJ (F), and AFD (G) neurons in both sexes using the *ins-39*(*syb4915)* reporter. n = 10-14 animals per group. Error bars are standard error of the mean (SEM). We performed a Mann-Whitney test in C, E-J, **** p < 0.0001, ** p < 0.01, * p < 0.05.

39 was expressed in five sets of neurons, AFD, ASK, ASJ, ASE, and AWC, all of which are ciliated sensory neurons with thermosensory, chemosensory, olfactory, electrosensory, and photosensory functions (Figure 4D). Using fluorescent dye staining of head ciliated sensory cells and AFD marker, we validated the higher GFP intensity specifically in ASK, ASJ, and AFD neurons in adult males compared to hermaphrodites (Figure 4E-G, Figure S6A-B). Taken together, INS-39 robust expression in sensory neurons in males from the earliest stages of development suggests unidentified male-specific functions.

### TRA-1 is necessary but not sufficient for *ins-39* dimorphic expression pattern

Does the genetic sexual identity of the animal determine *ins-39* dimorphic expression? In *C. elegans,* sexual differentiation that gives rise to dimorphic features is controlled autonomously in every cell by the activity of the TRA-1 master regulator^90^. Of note, neuronal *ins-39* expression precedes that of *tra-1*, suggesting an unknown, *tra-1-*independent regulatory mechanism for early sex-specific gene expression^91^. We first analyzed the *ins-39* locus for potential binding sites for TRA-1. We indeed identified a TRA-1 binding site in the *ins-39* first exon (Figure 5A). To investigate whether INS-39 dimorphic expression is determined by TRA-1 we manipulated its expression by masculinizing or feminizing the entire nervous system and scoring the number of neurons that express INS-39. Animals with a sex-reversed nervous system were generated by pan-neuronal expression of the *fem-3* gene (masculinization)^92^ or *tra-2(IC)* transgene (feminization)^93^. We found that masculinization of the nervous system significantly increased the number of INS-39-expressing neurons in hermaphrodites (Figure 5B-C). Conversely, pan-neuronal feminization decreased the number of INS-39-expressing neurons (Figure 5G-H) but was insufficient to reduce INS-39 to that of wild-type hermaphrodites (expression in AFD and sometimes in ASK). These results suggest that TRA-1 functions to restrict INS-39 expression in hermaphrodites, but additional factors are required for its dimorphic male expression.

**Figure 5:**
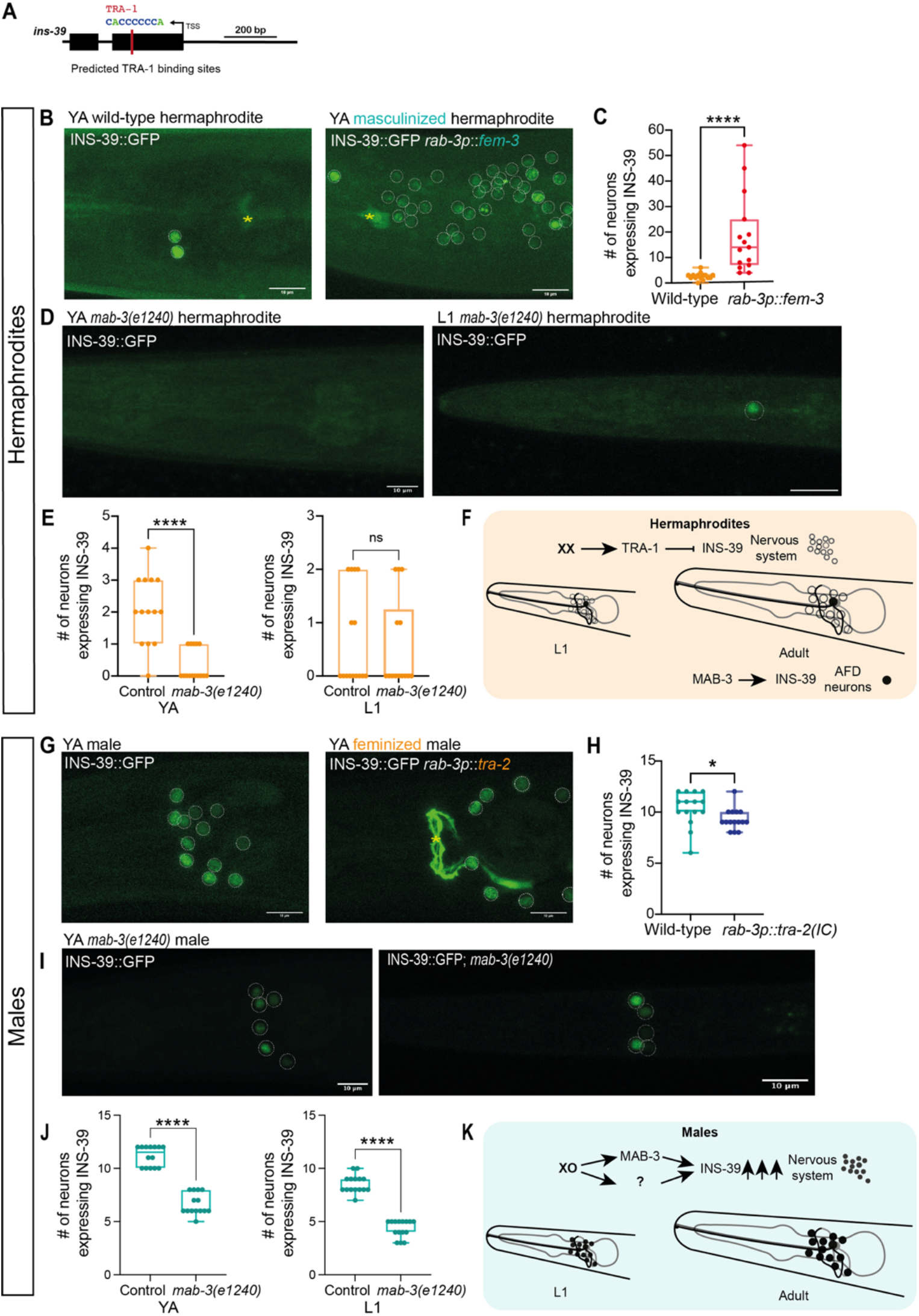
Neuronal sexual identity influences INS-39 spatial localization. (A) Schematic representation of predicted TRA-1 transcription factor binding site on the exonic region of *ins-39*. Binding-site was determined using RSAT matrix-scan search tool (see methods). (B) Representative confocal micrographs of the *ins-39*(*syb4915)* reporter expression in a wild-type hermaphrodite and pan-neuronally masculinized hermaphrodites, expressing *rab-3p::fem-3*. Co- injection marker is indicated by a yellow asterisk mark. Scale bars represent 10µm. (C) Quantification of the number of INS-39::GFP-expressing neurons from B. n=15 worms per group. (D) Representative confocal micrographs of the *ins-39(syb4915)* reporter expression in a wild-type YA hermaphrodite and *mab-3(e1240)* YA hermaphrodite. Scale bars represent 10µm. (E) Quantification of the number of INS-39::GFP-expressing neurons from wild-type YA (left panel) and L1 (right panel) hermaphrodite and *mab-3(e1240)* hermaphrodite. n=13-15 worms per group. (F) A model depicting the molecular elements mediating INS-39 expression in hermaphrodite at L1 and YA stage. (G) Representative confocal micrographs of the *ins-39*(*syb4915)* reporter expression in a wild-type male and pan-neuronally feminized male, expressing *rab-3p*::*tra-2(IC).* Co-injection marker is indicated by a yellow asterisk mark. Scale bars represent 10µm. (H) Quantification of the number of INS-39::GFP-expressing neurons from G. n=15 worms per group. (I) Representative confocal micrographs of the *ins-39(syb4915)* reporter expression in a wild-type YA male and *mab-3(e1240)* YA male. Scale bars represent 10µm. (J) Quantification of the number of INS-39::GFP-expressing neurons from wild-type YA (left panel) and L1 (right panel) male and *mab-3(e1240)* male. n=13-15 worms per group. (F) A model depicting the molecular elements mediating INS-39 expression in male at L1 and YA stages. Vertical bars in the box-and-whiskers graph (C, E, H, J) represent the median with dots showing all points from min to max. We performed a Mann-Whitney test for each comparison, **** p < 0.0001, * p < 0.05, ns-not significant.

The DMDs are among the known TRA-1 targets^94^. Screening previously published single- cell/bulk-sorted neuronal transcriptomic data sets for DMD genes expression that coincide with *ins-39* expression pattern^95–97^, we found that *dmd-9* is highly expressed in hermaphrodites in *ins- 39* expressing neurons (Figure S6C). Consistent with this, *dmd-9* mutant animals showed reduced expression of INS-39 GFP reporter in both sexes compared to wild-type animals (Figure S6D-E). *dmd-8* mutant animals did not show any significant change from wild-type animals, in line with its weak expression in INS-39-expressing neurons (Figure S6C-E). Given the interesting expression dynamics that our dataset captured for *mab-3*, with a strong peak at sexual maturation in both sexes, yet higher in males (Figure 2D), we decided to examine whether *mab-3* may have other sex-specific roles that have not yet been identified. We analyzed *mab-3* young adult hermaphrodite mutants and found that the number of INS-39::GFP-expressing neurons was significantly reduced (Figure 5D, E, left panel). Interestingly, *mab-3* loss of function did not affect L1 hermaphrodite INS-39 expression in AFD neurons (Figure 5E, right panel), again suggesting that other factors act early during development to regulate INS-39 levels. In males, *mab-3* mutations significantly reduced the number of INS-39::GFP-expressing neurons already early during development and sustained throughout adulthood (Figure 5I-J). Taken together, these results paint a complex and sexually-dimorphic regulatory path for INS-39 expression. In hermaphrodites, TRA-1 suppresses INS-39 expression throughout the nervous system, while *mab-3* antagonizes its function and promotes its expression in the AFD neurons (Figure 5F). In males, in the absence of TRA-1, *mab-3* and additional factors function to promote the extremely high *ins- 39* expression in many sensory neurons (Figure 5K). The emerging picture is of tight regulation of INS-39 in hermaphrodites, whereas in males, a more complex regulatory network coordinates high INS-39 neuronal expression.

### Sex-specific roles of INS-39 in survival and stress response in *C. elegans*

INS-39 is a member of an expanded class of insulin-like peptides in *C. elegans* and functions through an evolutionary conserved insulin-like growth factor signaling pathway^98^. All 40 insulin-like peptides identified in *C. elegans* are thought to act through a single receptor, DAF-2, the homolog of the human Insulin-like growth factor receptor IGFR1. Insulin/IGF1 signaling (IIS) regulates dauer entry, behavior, aging, development, and fat accumulation^99^. Recently, it was discovered that high temperature represses INS-39 expression in the AFD neuron of hermaphrodites^87^. We, therefore, examined the changes in INS-39::GFP expression in the AFD neurons of both sexes when subjected to temperature shifts (25°C and 15°C) from the cultivation temperature (22°C). As expected, we observed a significant reduction in INS-39 levels in AFD in hermaphrodites upon temperature shifts (Figure 6A-B). However, INS-39 levels in AFD in males did not change with temperature shifts and remained high (Figure 6A-C). We next tested INS-39 involvement in thermotaxis but found no apparent behavioral role for INS-39 in both sexes (Figure S7A-B). Considering the expression of INS-39 in ASJ neurons, which are known to be associated with cold tolerance, we reasoned that perhaps INS-39 might be involved in adult survival, a mechanism regulated by the IIS pathway^100^, and assayed *ins-39* mutants for cold tolerance. As previously shown^100^, hermaphrodites habituated to cooler temperatures (15°C) survived better during prolonged exposure to cold temperatures (2°C), in comparison to those that were grown at 22°C (Figure 6D). We found two dimorphic phenotypes for cold tolerance: (i) wild-type males survived chronic cold temperatures significantly better than hermaphrodites. This novel dimorphic phenotype, however, was *ins-39* independent as mutant males were indistinguishable from wild-type males (Figure 6D, left panel). (ii) cold-habituated hermaphrodites, but not males, showed improved survival rates in cool temperatures in an *ins-39-*dependent manner (Figure 6D, right panel). These results suggest a nuanced role of *ins-39* in cold survival.

**Figure 6:**
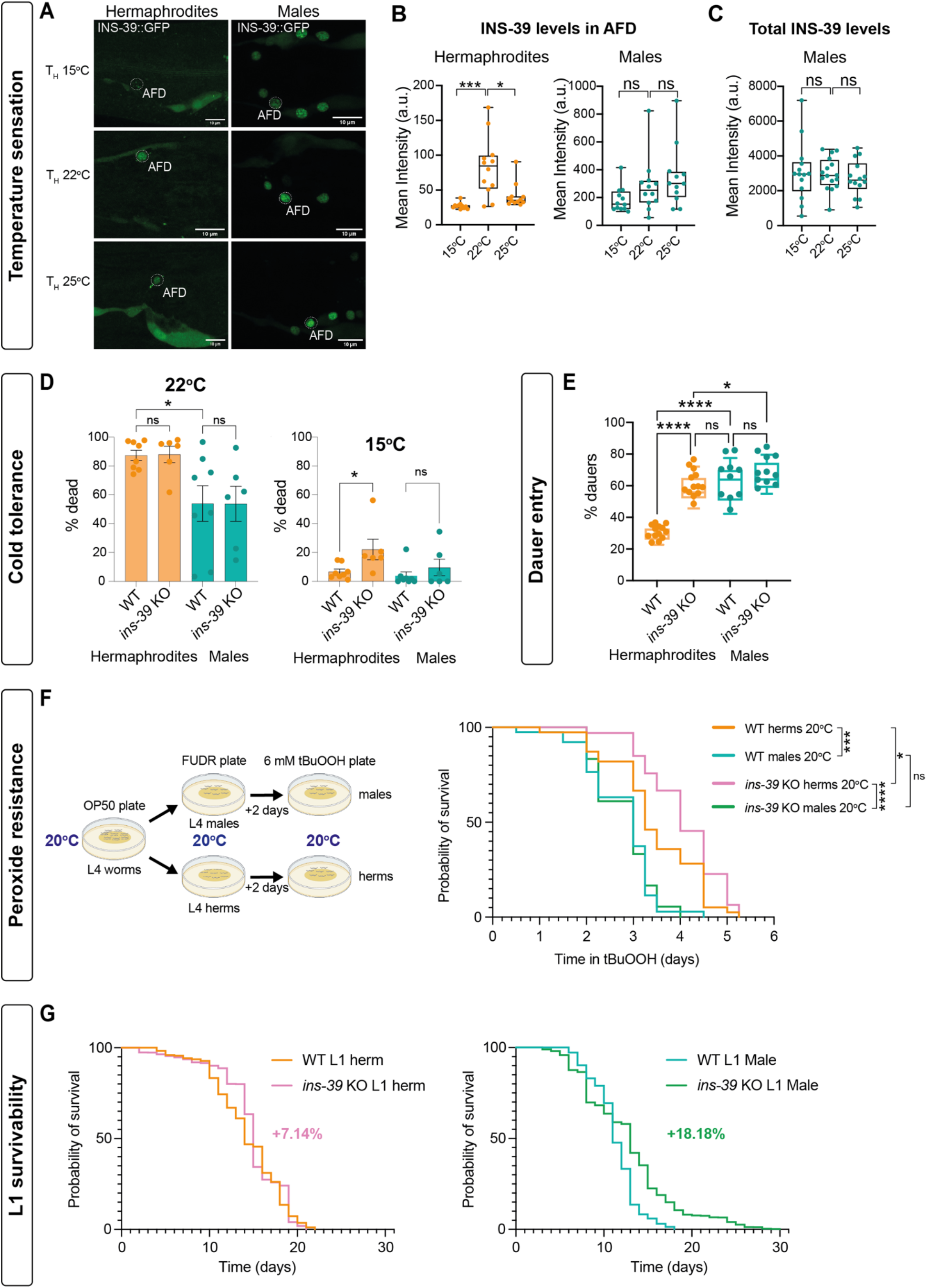
Functional characterization of the role of INS-39. (A) Representative confocal micrographs of the *ins-39*(*syb4915)* reporter expression in worms grown at 22°C and then transferred to 25°C (bottom) or 15°C (top) for 6 hours. Scale bars represents 10µm. (B) Quantification of mean GFP intensity from A in AFD in hermaphrodites (left) and males (right). n =11-12 worms per group. (C) Quantification of mean GFP intensity from A in INS-39 expressing neurons in males. n =12 worms per group. (D) Quantification of L1 lethality (cold tolerance) in wild-type and INS-39 KO CRISPR strain in hermaphrodites and males habituated at 22°C and 15°C for 6 hours. Each dot represents independent biological repeats. Error bars are standard errors of the mean (SEM). (E) Quantification of dauers percent in wild-type and INS-39 KO CRISPR strain in hermaphrodites and males habituated at 27°C (see methods). Each dot represents independent biological repeats. (F) Schematic of the survival assay on 6mM tBuOOH (left) and cumulative survival graph on 6mM tBuOOH of wild-type and INS-39 KO hermaphrodites and males (right). n =40 worms per group. (G) Cumulative L1 survival graph of wild-type and INS-39 KO hermaphrodites (left) and males (right) (see methods for n). Vertical bars in box-and-whiskers graph (B, C, E) represent the median with dots showing all points from min to max. We performed (in B, C, E) a Mann-Whitney test, (in D) a Dunnett’s multiple test for each comparison. The statistical significance for survival was calculated using the Log-rank (Mantel-Cox) test. **** p < 0.0001, *** p < 0.001, ** p < 0.01, * p < 0.05, ns-non- significant.

*ins-39* was previously shown to play a role in nematode survival against peroxide stress through the insulin signaling pathway^87^. We confirmed the previously reported enhanced peroxide survival in INS-39 mutant hermaphrodites grown and assayed at 20°C or at 25°C (Figure 6F, Figure S7C- F). However, unlike hermaphrodites, our results show that *ins-39* does not play a significant role in males, regardless of pre-exposure to higher (25°C) or lower (20°C) temperatures (Figure 6F, Figure S7C-F). Furthermore, we observed that wild-type males exhibited lower survival rates against peroxide compared to hermaphrodites, although this phenotype was not dependent on INS- 39 expression in males (Figure 6F).

Since *ins-39* levels are much higher in males already early during development, we reasoned this might give them some advantage over hermaphrodites in coping with early life stress. Therefore, we focused on early life stress using two strategies. We first tested the ability of the animals to enter the dauer stage and found that wild-type males were more efficient in dauer transition than hermaphrodites (Figure 6E). Unexpectedly, hermaphrodites mutant for *ins-39* became as efficient as wild-type males in dauer entry, while *ins-39* mutant males were similar to wild-type males (Figure 6E), suggesting that *ins-39* expression has a dimorphic role in the dauer pathway.

Lastly, we tested L1 survivability. The IIS pathway has been shown to regulate L1 arrest, in which worms can cease growth and development as young larvae in the absence of food^101^. The survival of L1-arrested animals can be shortened or lengthened depending on the degree of IIS activity. With this in mind, we evaluated if *ins-39* had any sex-specific regulation of L1 survival. We first found that wild-type male worms exhibited a significantly lower survival rate (21.42% shorter) as arrested L1 larvae than hermaphrodites (Figure S7G). Interestingly, *ins-39* KO L1 animals of both sexes survived significantly longer than their wild-type controls, and males even more so than hermaphrodites (Figure 6G). These results suggest a negative role for *ins-39* in the regulation of L1 survival, with a more critical role in males than hermaphrodites.

Taken together, we suggest that the lower expression levels of INS-39 in hermaphrodites is fine-tuned to critically respond to environmental changes, providing a survival advantage in the face of unfavorable conditions. In contrast, the higher expression in males might hinder its functionality, creating a ceiling effect.

## DISCUSSION

Our transcriptomic atlas provides a much-needed molecular roadmap with which important developmental and molecular questions can be answered in a dimorphic context. We report a comprehensive gene expression atlas for both sexes of *C. elegans* throughout development which has been lacking until now. Our findings revealed 519, 63, 614, 2141 and 3564 genes that were differentially/dimorphically expressed in L1, L2, L3, L4 and YA, respectively, providing a more comprehensive sex and stage datasets compared to previous findings^27, 28, 46, 102, 103^. The observed increase (∼12 times more) in the amount of identified dimorphic genes compared to previous studies^28^ can be attributed to restrictions of using pseudo males to compare between stages in both sexes, resulting in a lower readout. Importantly, we established and refined a novel protocol to isolate healthy males from the very first larval stage of development for the benefit of the *C. elegans* research community.

Across different organisms, the expression levels of most genes change during development. In addition, sex bias in gene expression can vary greatly between and within species and depends on factors like species, tissue type, RNA-seq library preparation, sample size, and statistical criteria ^104–106^. Several novel observations can be made when sex-enriched mRNAs at various stages are compared. It appears that sexual differentiation is first manifested by a small subset of sex-enriched genes in early larvae and a larger, more specific subset of mRNAs in later larvae throughout all developmental stages. Gene ontology enrichment of TFs during the pre-sexual maturation period showed a substantial enrichment for processes essential for promoting subsequent dimorphic development, as well as sexually dimorphic morphogenic processes. These TFs are likely to contribute to or drive the development of sex-specific traits and structures, which emerge later during sexual maturation. Additionally, there are more male-biased DEGs than hermaphrodite-biased DEGs. This male bias in differential gene expression has been observed in other organisms as well, ranging from plants to insects^107–109^. In humans, sex-biased gene expression is largely tissue-specific, and sex-biased genes exhibit nonrandom and tissue-specific genomic distribution^10^.

Although the development of males and hermaphrodites is similar before sexual maturation, several crucial male cell fates are already determined at hatching. For example, three significant groups of male-specific blast cells give rise to the male mating structures^110^. Our list of male-enriched genes could point to additional genes that are involved in the regulation of male identity. The almost complete male-biased expression of neuronal genes could be explained by the size of the male nervous system, which contains 30 percent more neurons, as well as a larger connectivity network of chemical and electrical synapses. The unique neuro-peptidergic blueprint of males is a novel outcome of this study. We find that neuropeptide coding genes are differentially expressed between the sexes and across development, with a strong bias towards higher male expression.

In rodents, Neuropeptide Y (NPY) expression in many brain areas under basal, unstressed conditions is lower in females than in males^111, 112^. We show similar results for the insulin peptide *ins-39*, which exhibits strikingly higher and broader expression levels in males versus hermaphrodites but, puzzlingly, is necessary almost exclusively in hermaphrodites for sensing and coping with various harsh environmental conditions and stressors. For the NPY system, it has been speculated that the lower expression levels might put females at a disadvantage in dealing with stress^112^. Our opposite results for *ins-39* prompt an alternative hypothesis, in which the low expression levels provide a better environmental endurance or read out, making the animal more fine-tuned to its surroundings. This hypothesis is further substantiated by the tight regulation of *ins-39* expression levels in hermaphrodites. Why males require high *ins-39* levels is unknown but may represent an evolutionary cost that comes as a tradeoff for an unknown advantage that awaits further research.

Interestingly, we found that the highest number of common DEGs was shared between L1 and YA. This perhaps points to these stages as critical windows of sexually dimorphic development, providing the needed flexibility to respond to environmental cues, a phenomenon observed in memory imprinting and early-life experience studies^113^.

Very few DEGs in our dataset were common to all development stages. Out of the 769 TFs described so far in *C. elegans*^53^, we identified a total of 218 TFs with temporal and sexual specificity throughout development. Although previous research^27^ identified 17 TFs, they were constrained by sample size, thus our data provides better temporal resolution of transcription factors in either sex, as exemplified by *ces-2*, which was consistently greater in males across all developmental stages. *ces-2* controls asymmetric cell division and mitotic spindle orientation in the neurosecretory motor (NSM) neurons lineage and is necessary to initiate programmed cell death^114, 115^. Our study predicts that *ces-2* could be involved in establishing a male cell identity from the very first larval stage, therefore crucial for further research.

Although our approach presents the most comprehensive gene expression dataset of dimorphic development to date, this study is not full, for it was not designed to identify transcripts in embryonic stages^116^, post-transcriptional regulation of genes, or dimorphic effects elicited by hormones as in mammals^117^. Nevertheless, this work does not only enrich the *C. elegans* community with new methodologies and expression data, but also offers 1047 new conserved dimorphic candidate genes with their associated human genetic disorders and traits that may underlie the sexually-dimorphic molecular characteristics of various human diseases.

## Supporting information

strains and reagents

## ACKNOWLEDGEMENTS

We thank members of the Oren-Suissa lab for their critical insights regarding the manuscript. We thank Dr. Shifra Ben-Dor at Bioinformatics Unit, Life Sciences Core Facilities of Weizmann Institute of Science for helping with promotor analysis. RNA-seq library preparation was done with critical advice from Dr. Hadas Keren-Shaul, Dr. Merav Kedmi and Dr. David Pilzer from the Genomics Sandbox unit at the Life Science Core Facility of Weizmann Institute of Science. We are grateful to Patrick Laurent for sharing strains. We thank Ronen Hayun, Maayan Maron, Eli Hotoveli, Anastasia Zarankin, Avigail Gedanken, Yarden Tala and Moshiko Hafzadi at Design & Development of Weizmann Institute of Science for the website development. Some strains used in this study were obtained from Caenorhabditis Genetics Center (CGC), which is funded by the NIH Office of Research Infrastructure Programs (P40 OD010440). We thank WormBase, an online biological database for *C. elegans*, which is supported by Grant U41 HG002223 from the National Human Genome Research Institute at the NIH, the UK Medical Research Council, and the UK Biotechnology and Biological Sciences Research Council. MOS acknowledges financial support from the European Research Council ERC-2019-STG 850784, Israel Science Foundation grant 961/21, Dr. Barry Sherman Institute for Medicinal Chemistry, Sagol Weizmann-MIT Bridge Program and the Azrieli Foundation. MOS is the incumbent of the Jenna and Julia Birnbach Family Career Development Chair. RH is greatly thankful to Council of Higher Education, Israel for PBC postdoctoral fellowship.

## AUTHOR CONTRIBUTIONS

R.H. conducted and analyzed the experiments. H.S. contributed to the imaging, crossing and analysis of INS-39 expression. E.L. helped in imaging NeuroPAL strains. Y.S. conducted CRISPR/Cas9-mediated genome editing. S.K. conducted thermotaxis assays. S.K. and Y.S. carried out cold tolerance assay. G.S. and H.G. carried out the bioinformatics analysis, under the supervision of O.R and M.O.S. M.O.S. supervised and designed the experiments. R.H., Y.S. and M.O.S. wrote the paper.

## RESOURCE AVAILABILITY

### Lead Contact

Further information and requests for resources and reagents should be directed to and will be fulfilled by the Lead Contact, Meital Oren-Suissa.

### Material Availability

Unique strains generated in this study have been deposited at the Caenorhabditis Genetics Center. Requests for other strains and plasmids should be directed to the Lead Contact.

## MATERIALS AND METHODS

### *C. elegans* strains and maintenance

All *C. elegans* strains were cultivated as per standard methods^29^. Wild-type strains were Bristol, N2. *him-5(e1490)* or *him-8(e1489)* were treated as controls for strains with these alleles in their background. Worms were grown on nematode growth media (NGM) plates seeded with *E. coli* OP50 bacteria as a food source. Worms were raised at 20°C, unless noted otherwise. The sex and age of the animals used in each experiment are noted in the associated figures and legends. All transgenic strains used in this study are listed in key resources table.

### Generation of *C. elegans* male cultures

To generate pure male populations, temperature-sensitive mutation (*y1*) in the dosage compensation gene, *dpy-28* along with *him-8(e1489)* mutations were employed^118^. *dpy-28(y1) him-8(e1489)* worms were cultured at 15 °C in 15 cm NGM plates seeded with OP50. Embryos were isolated by hypochlorite treatment of gravid adults collected from 3-4 NGM plates. The obtained embryos were hatched for 14-16 hours on a foodless NGM plate at a restrictive temperature 25°C. At restrictive temperature, most hermaphrodites die as embryos or L1 larvae, whereas the XO males are unaffected. After 14-16 hours, L1 larvae and unhatched embryos were collected in 2 ml M9 buffer and pelleted at 3000 rpm for 1 min. Approx. 5000 L1/embryos were placed onto side (area without food) about 1.5 cm from the food on a food NGM plates (Figure S1). Depending upon the density of the pellet the number of plates were increased accordingly. Plates were incubated at RT for 1-2 hours allowing the L1 animal to crawl towards food. After, incubation the area were L1/embryos were placed were cut out using scalpel. L1 worms which were on food were collected using 1 ml M9 buffer and washed 5 times with M9 buffer to get rid of bacteria. The L1 animals were counted and about 25,000 worms were placed on 15 cm NGM plates with OP50 at 25°C until they reached their maturity or to a desired time point. The percentage of males obtained were counted for each set of experiments. A schematic representation of this protocol is illustrated in Figure S1.

### Automated worm tracking

Day 1 adult *him-8(e1489)* males and *dpy-28(y1) him-8(e1489)* males were assayed for their speed/locomotion and body area. Adult males from both groups were placed onto a NGM plate seeded with 30 μL of OP50 bacteria. 2-4 worms were placed onto food and a 1.5 cm diameter plastic was placed around them so that they do not crawl away from the camera’s field of view. The plate was placed inside tracker and worms were left to habituate for 10 min. Using WormLab automated tracking system (MBF Bio-science)^119^, the worms were recorded for 2 min at RT. To recover the worm contour and skeleton for phenotypic analysis, the collected videos were segmented, and body area, speed, track length, mean amplitude, cumulative reversal time, and cumulative forward time metrics were exported. Prism (GraphPad) was used for statistical analysis.

### Body length and width measurement

Day 1 adult *him-8(e1489)* males and *dpy-28(y1) him-8(e1489)* males were imaged in DIC channel using Zeiss LSM 880 confocal microscope with 20x objective lens. The images were imported to ImageJ/Fiji software, version 2.3.0/1.53q, and a line was drawn from the tip to tail, or across the width of each worm. The length of the line was measured and was statistically analyzed.

### Mating behavior assay

Mating assays were conducted based on previously established methods^22, 120^. 15-20 early L4 males from the test group and 15–20 early L4 hermaphrodites from *unc-31(e928)* group were separated from a mix stage population and were transferred to separate fresh OP50 NGM plates and incubated at 20°C until they reached sexual maturation. On a fresh plate seeded with a thin lawn of OP50 bacteria, 10 adult *unc-31(e928)* hermaphrodites were transferred. *unc-31(e928)* hermaphrodites do not move a lot, allowing for an easy capturing of male behavior. A single virgin male from the test group was added to this lawn. Animals were kept under observation, and the course of events was recorded using a Zeiss Axiocam ERc 5s mounted on a Zeiss stemi 508 for a period of 15 minutes, or until the male ejaculated, whichever occurred first. Mated males and hermaphrodites were discarded from the plate after the event was completed. A new hermaphrodite was then added to the lawn to keep the number of hermaphrodites same between experiments. The video was analyzed, and the males were scored for their time until successful mating, contact response and vulva location efficiency. Males that did not mate within 15 minutes time window were not analyzed. Contact responses were defined as mail tail apposition and start of backward movement on the hermaphrodite’s body. Percentage response to contact was calculated using the formula (the number of times a male showed contact response/the number of times the male contacts a hermaphrodite via the rays * 100^22, 120^. Vulva location efficiency (L.E.)^121^, was calculated using formula 1/number of major passes or hesitations at the vulva until ejaculation or 15 min time window.

### RNA Isolation, and library preparation

More than 2000 worms from each developmental stage and sex were collected from agar plates and were washed 5 times using M9 buffer to get rid of bacteria. Each set contained 4 biological replicates. Worms pellet were collected in 500 µL of Trizol and flash frozen in liquid nitrogen and stored at -80°C until further processing. Total RNA was prepared from the frozen worm pellets following the instructions for the TRIzol LS (Invitrogen) protocol. After the isopropanol precipitation step, the RNA was re-suspended in the extraction buffer of the RNA isolation kit (PicoPure, Arcturus), and further isolation was carried out in accordance with the manufacturer’s instructions. When starting with a smaller number of worms, this two-step purification technique aids in getting RNA of good quality. RNA concentration was determined by Qubit Fluorometer (Invitrogen, Model #4), and RNA integrity was checked using Bioanalyzer (Agilent). Only samples showing RIN^e^ ≥ 8 was further processed. To create RNA-seq libraries for the expression profiling, a bulk version of the MARS-Seq procedure^40, 41^ was employed. Briefly, reverse transcription was used to barcode and pool 18 ng of input RNA from each sample. The pooled samples underwent second strand synthesis after Agencourt Ampure XP beads cleanup (Beckman Coulter), and they were linearly amplified by T7 in vitro transcription. The resulting RNA was fragmented and then converted into final library by tagging the samples with Illumina sequences during ligation, RT, and PCR. Finally, libraries were quantified by Qubit and TapeStation followed by qPCR for *lmn-1* gene as previously described^40, 41^. Sequencing was carried out using an Illumina Nextseq 75 cycles high output kit with paired end sequencing. Median, mean and standard deviation of the sequencing depth of all samples were 14.08, 15.02, 5.38 million reads.

### RNA-seq data analysis

Raw next-generation sequencing (NGS) data in the FASTQ format were analyzed using user-friendly transcriptome analysis pipeline (UTAP)^122^. Briefly, reads were trimmed using “cutadapt”^123^ and mapped to the *C. elegans* reference genome WBcel235 using STAR v2.4.2a^124^. UMI counting was done using HTSeq-count^125^ after marking duplicates in union mode. Differential gene expression analysis and normalization of the counts was performed using DESeq2^77^ using the rld (log2 normalized) gene expression values on a per-developmental stage basis between sexes. Raw p-values were adjusted using the Benjamini and Hochberg method^126^. A threshold of p-adjusted ≤ 0.05, |log2FoldChange| ≥ 1 and baseMean >= 5 was used to identify genes with significant differential expression. The WormBase gene set enrichment analysis tool was used for functional annotation of gene ontology terms that are over-represented in differentially expressed genes for each developmental stage and sex^50, 85^. A q-value threshold of 0.1 was used for every analysis. Jvenn^49^ was used to create diagrams.

### Gene conservation analysis between *C. elegans* and Human

Human orthologs for the *C. elegans* genes and their associated Online Mendelian Inheritance in Man (OMIM) human-disease phenotypes were downloaded from OrthoList 2 (OL2)^127^, a compendium of *C. elegans* genes with likely human orthologs (http://ortholist.shaye-lab.org/). The genes from our RNA-seq datasets were overlapped with the list from OrthoList 2 and were incorporated into analysis.

### Microscopy

Worms were anaesthetized on a drop of 200 mM sodium azide (NaN3, prepared in M9 buffer) mounted on a freshly prepared 5 % agarose pad on a glass slide. Images were acquired using Zeiss LSM 880 confocal microscope with 63x objective lens unless otherwise noted and processed using ImageJ/Fiji software version 2.3.0/1.53q^128^. All images of the same group were imaged using identical microscope settings. For expression of reporters, representative images with maximum intensity projections are shown for all channels. Figures were prepared using Adobe Illustrator v24.0. For imaging NeuroPAL worms, channels were pseudo-colored in accordance with^129^.

### BioSorter analysis

*C. elegans* strains *ins-39p::gfp* or *him-5* (control) were grown and maintained at 20°C. L1 synchronized nematode cultures were obtained by standard bleaching protocols. Eggs were allowed to hatch overnight on foodless (i.e., without a bacterial lawn) 60 mm NGM plates, then L1-arrested larvae were collected in 50 ml conical tubes and were resuspended in S-basal at a final density of about 2000 worms/ml. Worms sorting was performed on a BioSorter equipped with a 250 FOCA and S-basal as the sheath fluid. To isolate L1 males, *ins-39p::gfp* strain was used (GFP positive in head neurons). Non-fluorescent *him-5* strain was used to set gates for GFP positive worms and to exclude auto-fluorescent worms. Sorted worms were collected onto a 60 mm OP50 plate.

### Neuron identification using NeuroPAL

For imaging NeuroPAL worms, the protocol followed are in accordance with^129^. For INS-39 CRISPR GFP neuronal identity, colocalization with the NeuroPAL landmark strain OH15262 harboring *otIs669* transgene was used to determine the identity of all neuronal expression.

### DiO/DiD staining

Age synchronized worms were washed thrice with 500 μl M9 buffer in 1.5 ml microfuge tube and incubated in dark for 1 h in 1 ml M9 containing 5 µl DiD dye (Vybrant™ DiD Cell-Labeling Solution, ThermoFisher) at ∼20-30 rpm. The worms were then centrifuged, and worm pellet was transferred to a fresh plate and animals were let to move onto the bacterial lawn.

### RT-PCR

Total RNA was extracted from 10 to 20 µl of age-synchronized packed worms’ pellet (washed three times with M9 buffer) from relevant genotypes/developmental stage using protocol described above under “RNA Isolation, and library preparation” header. The isolated total RNA was then converted to cDNA using SuperScript™ IV first-strand synthesis system (Invitrogen, Catalog # 18091050) with random hexamers. The concentration of cDNA was determined using NanoDrop microvolume spectrophotometers and fluorometer. qPCR reactions were set up in 10 µl volume using Fast SYBR™ Green Master Mix (Applied Biosystems, Catalog # 4385612) as per manufacturer protocol. Primers used are indicated in key resources table. Ct values were retrieved and relative quantification of the expression of the target genes was performed utilizing 2−ΔΔCt method.

### CRISPR/Cas9-mediated genome editing

For creation of *ins-39(ety9)* deletion strain, previously described CRISPR/Cas9 genome engineering protocol was employed^130^. Briefly, tracrRNA and three crRNAs targeting the *dpy-10* locus and the boundaries of the *ins-39* locus, were combined with a recombinant cas9 (IDT) and supplemented with ssODN repair template to introduce a dominant mutation into the *dpy-10* locus^130^ and a second ssODN composed of the 5’ and 3’ flanking sequences of the *ins-39* locus boundaries (see key resources table for sequences). Rol/dpy F1 progeny were singled and screened by PCR using the primers ATAGCAGAACATGGGCATCC and AACCGTTGGGTATTTGACCA, that amplify a 735bp product only in *ins-39* deleted animals. Plates with a correct PCR signal were homozygozed and Sanger-sequenced to validate the accuracy of the *ety9* edit. Resulting strain was backcrossed 2 times to lose the *dpy-10* mutation.

The *syb4915*, *syb4956* and *syb4992* allele was generated by SunyBiotech.

### Promotor analysis

*ins-39* promotor region was extracted from WormBase by taking 1000 bp upstream and 500 bp downstream of the transcription start site which included the whole protein coding part of the gene. *tra-1* binding sites were derived into the matrix from previously published study^131^ using genomatix genome analyzer tool. The promotor region binding sites were determined by using RSAT matrix-scan search tool.

### Thermotaxis assay

Briefly, age-synchronized animals were sex-separated at late L4-YA stages and moved to 15°C, 22°C and 25°C for 6 hours. The animals were then washed and placed onto the thermotaxis plate.

A modified steep linear gradient was maintained on a 10cm plate using an aluminum plate with a hot plate (set at 27°C) on one side and an ice pack on the other^132^. Temperature on the agar surface was measured using a temperature gun. The gradient was allowed to stabilize for 5 mins, after which the animals were introduced at 22°C isotherm and allowed to roam freely for 20 minutes. The animals were killed by quickly transferring to -20°C for 5 minutes. Animals were scored for positive or negative thermotaxis as 1cm from the center of origin towards either side. Thermotaxis index was calculated for each experimental group as:

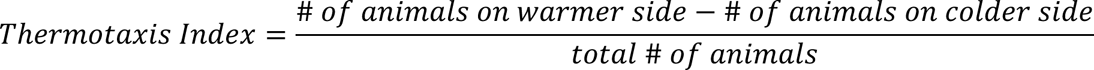

### Cold tolerance assay

Cold tolerance assay was conducted as previously described^100^. Briefly, age synchronized animals were sex-separated into males and hermaphrodites at late L4-YA stages. Then, they were moved to either 15°C or 22°C for 6 hours. After which, they were washed twice with M9 and plated onto OP50 plates and moved to 2°C for 48 hours and then scored for live/dead. The percentage dead were calculated from the total animals on the plate. All experiments were conducted in 4 independent biological trials.

### Dauer formation assay

Dauer formation assay induced by high temperatures was conducted following previous protocols^133^, with slight modifications. 50 gravid adults were collected that had been continuously raised at 15°C were placed on OP50 plates for 6 hours at room temperature. The hermaphrodites were removed, and the laid eggs were incubated at 27°C for 44 hours, while control plates were kept at room temperature for the same duration. To examine the presence of dauers (active, thrashing animals) and non-dauers (inactive, non-thrashing animals), we flooded the assay plates with 2 mL of 1% SDS solution, swirled them, and allowed them to incubate for 15 minutes at room temperature. The number of dauers and non-dauers animals were recorded and analyzed. All experiments were conducted in 3 independent biological trials.

### Survival assays in tert-butyl hydroperoxide

The survival assay was conducted on solid agar plates containing tert-butyl hydroperoxide at a final concentration of 6 mM, following previous protocols^87^, with slight modifications. To prepare the assay plates, tert-butyl hydroperoxide was added to molten agar maintained at 55°C, and 12 ml of the mixture was poured onto 60 mm NGM agar plates. The plates were left to dry at room temperature for 2 days and then seeded with 200 µl of a 5x concentrated OP50 culture that had been grown overnight, pelleted, and resuspended in M9. The worms were cultured on standard OP50 plates until they reached the L4 developmental stage and then separated by sex into groups of up to 100 on plates containing 10 μg/ml 5-fluoro-2′-deoxyuridine (FUDR). For each genotype and sex, 40 Day 2 adults were transferred to assay plates at the specified temperature. The worms were monitored every 6-12 hours and scored as alive, dead, or censored until all worms had died. A worm was considered “dead” if it did not show any visible movement in response to a gentle touch from a poking lash, while worms that were lost or had desiccated on the side of the plate were considered “censored.” Manual scoring was conducted, and the resulting data was plotted and analyzed. All experiments were conducted in 3 independent biological trials. To compare the survival rates Kaplan-Meier method was used. The statistical significance was calculated using the Log-rank (Mantel-Cox) test in Prism (GraphPad).

#### L1 survival assay

L1 survival assay was conducted as described previously with slight changes^101^. Briefly, L1 animals were obtained from a well-fed *C. elegans* plates from the indicated genotype employing the same method as described to isolate male under section “Generation of *C. elegans* male cultures” until L1 stage. This was done to keep all the parameters same across all the experimental group. The L1 worms were collected using M9 from the foodless plate and were diluted to a final concentration of no more than 10 L1 worms/µl in using M9. This was done because the survivability of *C. elegans* at L1 arrest depends on the density of worms^101^. Starvation cultures were placed in 40 ml glass vials and incubated on a rocker-shaker at 20°C at low-speed. Each day, aliquots containing at least 50 L1 arrested worms from each group were removed and plated on a foodless plate. The live/dead and total number of worms were recorded each day. All experiments were conducted in 3 independent biological trials. To compare the survival rates Kaplan-Meier method was used. The statistical significance was calculated using the Log-rank (Mantel-Cox) test in Prism (GraphPad).

### Quantification and statistical analysis

For specific analyses, information about quantification, statistical testing, sample size can be found in the figure legends and method details section.

### Transgenic Strains/Molecular cloning

To generate pan-neuronal masculinized or feminized strains *tra-2[intracellular]*^134^ or *fem-3* was expressed under *rab-3p*. See supplementary file SX for primers.

### Data availability

The RNA-seq datasets associated with this article are included in supplementary table S2. The sequence data of this article have been submitted to NCBI and are available under BioProject (TBA).

