## Supplemental figures for "Sex-Specific Developmental Gene Expression Atlas Unveils Dimorphic Gene Networks in *C. elegans*"

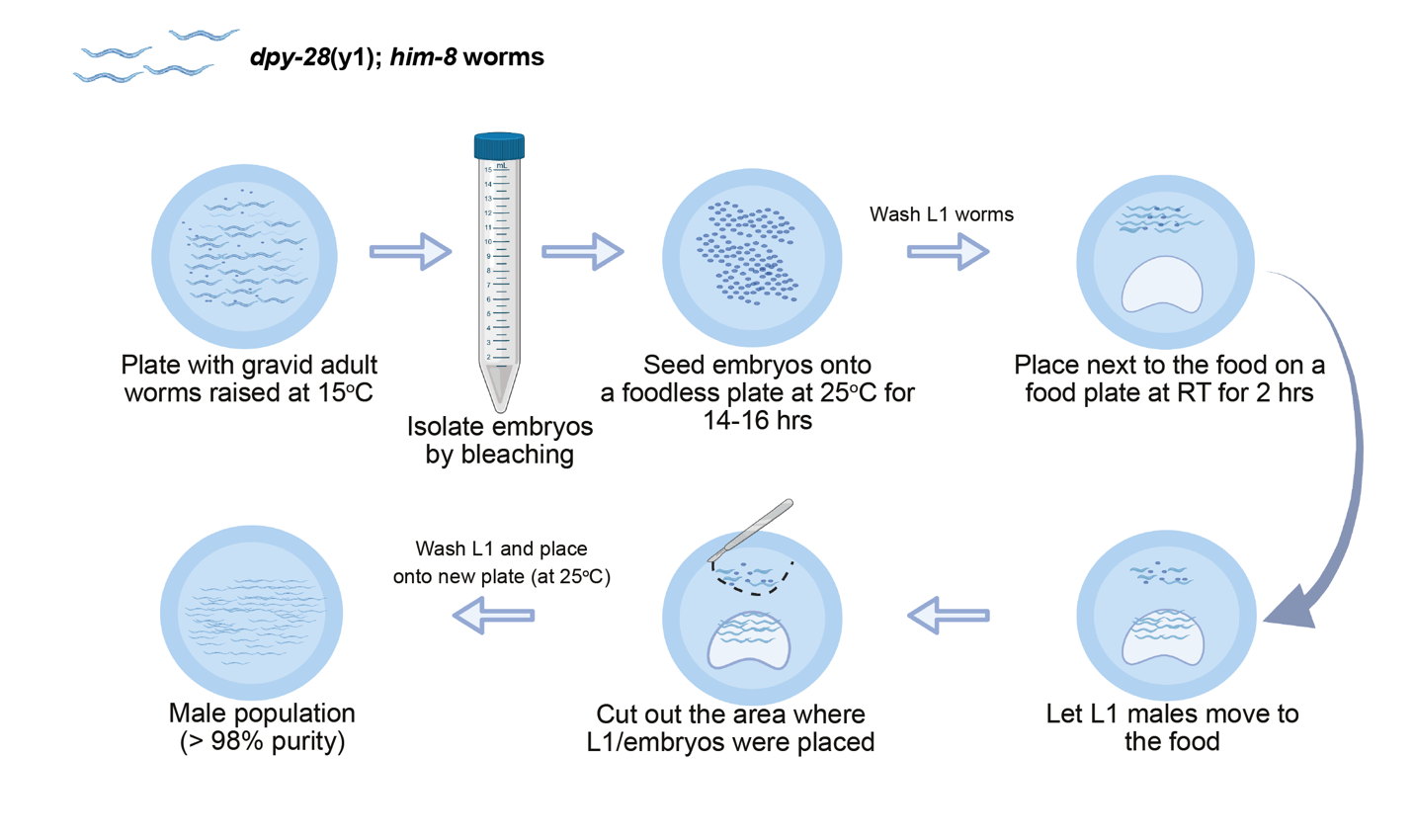

**Figure S1: Protocol for large scale male isolation in early larval stage of *C. elegans.***

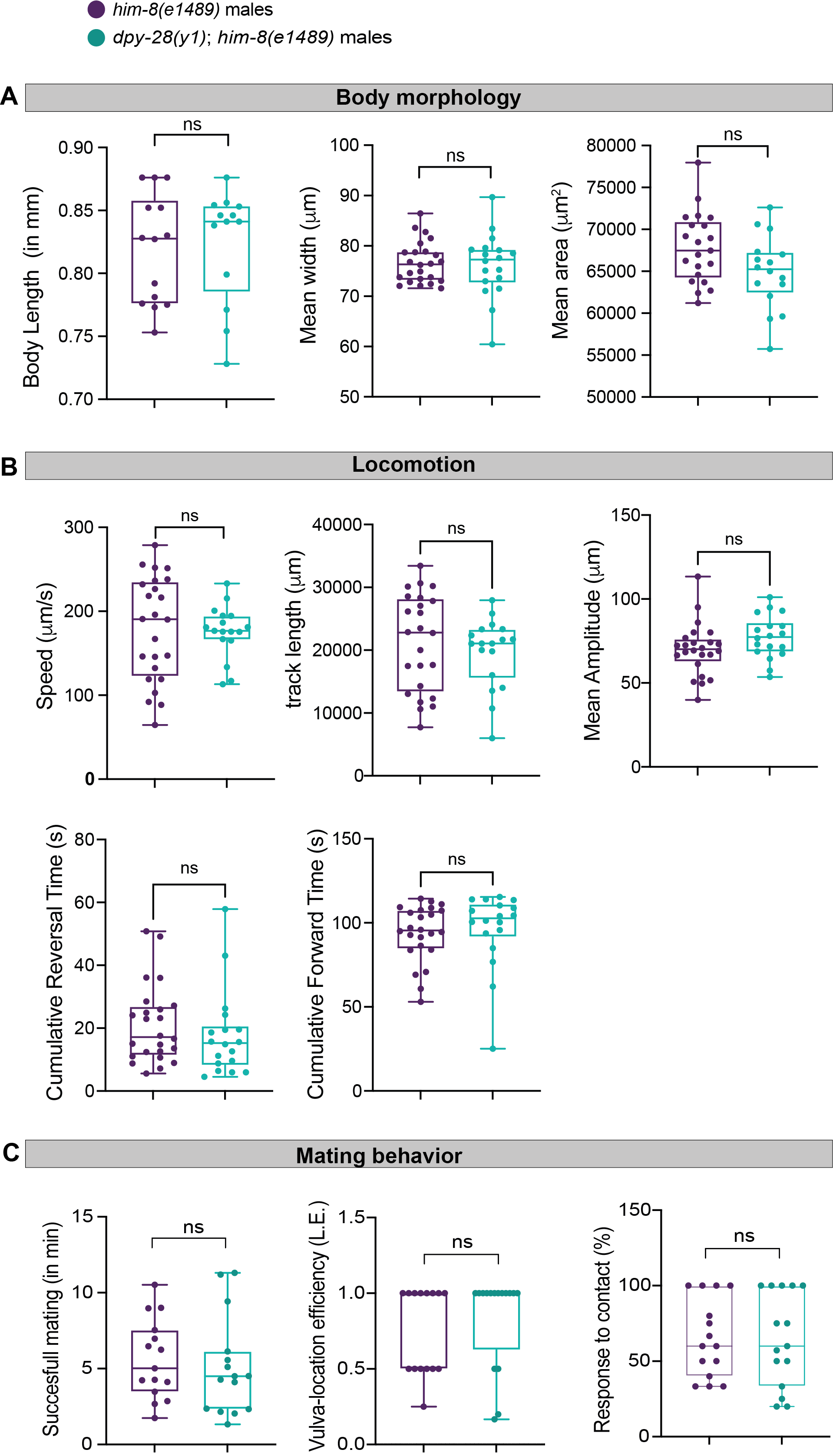

**Figure S2: *dpy-28* males are identical to wildtype males.**

**(**A) Quantification of body morphology. Body length (left), n = 13-14 worms per group, mean width (center), n = 18-24 worms per group, mean area (right), n = 16-21 worms per group.

(B) Quantification of locomotion. Speed (up-left), n = 17-24 worms per group, track length (up-center), n = 18-24 worms per group, mean amplitude (up-right), n = 18-24 worms per group, cumulative reversal time (down-left), n = 18-24 worms per group, cumulative forward time (up-center), n = 18-24 worms per group.

(C) Quantification of mating behavior. Time until first mating (left), vulva-location efficiency (center) and response to contact (right), n = 15-16 worms per group of *dpy-28*(*y1*);*him-8*(*e1489*) males and *him-8*(*e1489*) males.

Vertical bars in the box-and-whiskers graph represent the median, with dots showing all points from min to max. We performed a Mann-Whitney test for all comparisons, ns- non-significant.

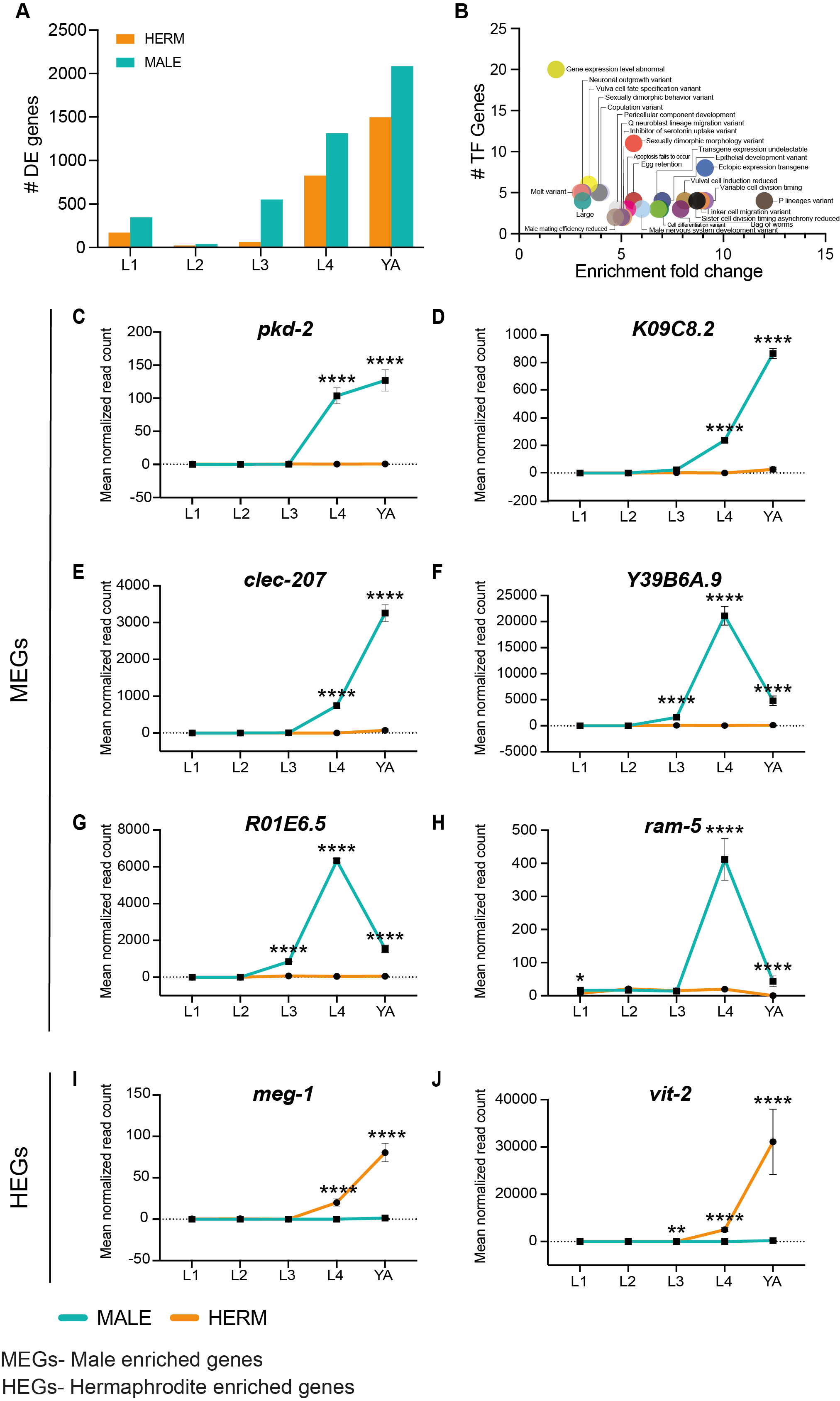

**Figure S3: Expression of known sex-specific genes rises during or after sexual maturation**.

(A) Bar graph representing number of differentially expressed genes across all the developmental stages in both the sexes (cyan: males and orange: hermaphrodites).

(B) Phenotype GO term enrichment analysis of differentially expressed transcriptional factors before sexual maturation (until L3). Each circle represent GO term. The significance of the enrichment was analyzed using q-value threshold of 0.1.

(C-J) Normalized read counts of *pkd-2* (C), *K09C8.2* (D), *clec-207* (E), *Y39B6A.9* (F), *R01E6.5* (G), *ram-5* (H) *meg-1* (I) *vit-2* (J) across all the developmental stages in both the sexes (cyan: males and orange: hermaphrodites). Error bars are standard error of the mean (SEM). adjusted p-values were calculated by Wald test for each comparison performed by DESeq2^1^, **** p < 0.0001, ** p < 0.01, * p < 0.05.

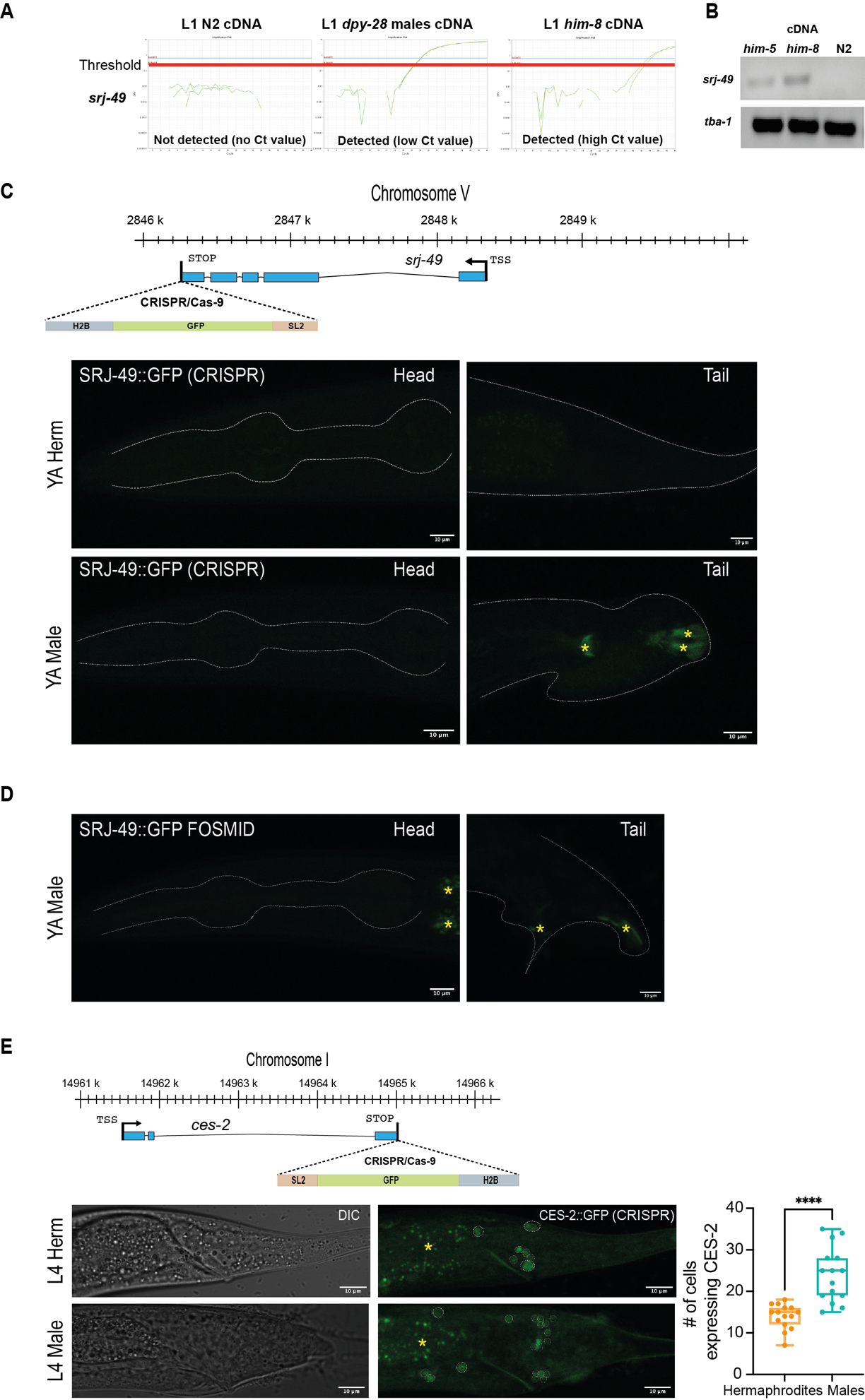

**Figure S4: *srj-49* mRNA is detectable only by RT-PCR only in *C. elegans* strains containing high male ratios.**

(A) Real-time qPCR amplification curve of *srj-49* mRNA at L1 stage of male-enriched strains (*him-8* and *dpy-28*(*y1*);*him-8* males) and wild-type N2 hermaphrodites-containing control strain of *C. elegans*. Each line (green) represent a biological repeat.

(B) DNA electrophoresis of PCR products amplified using *srj-49* and *tba-1* (tubulin) primers from cDNA isolated from mixed populations of *him-5*, *him-8* and N2 plates. *srj-49* mRNA is only detected in strains containing male populations and not detected in wildtype N2, that typically contains only hermaphrodites.

(C) Top, schematic of the CRISPR/Cas9 genome editing strategy used to engineer the *srj-49(syb4956)* CRISPR reporter. Bottom, representative confocal micrographs of head and tail of young adult hermaphrodite (top) and head and tail male (right) carrying the SRJ-49::GFP fosmid. Head pharynx and tail are outlined in white dashed line. Autofluorescence is indicated by yellow asterisk mark. Scale bars represents 10µm.

(D) Representative confocal micrographs of head and tail of young adult male right carrying the *srj-49(syb4956)* reporter expression. Head pharynx and tail are outlined in white dashed line. Autofluorescence is indicated by yellow asterisk mark. Scale bars represents 10µm.

(E) Top, schematic of the CRISPR/Cas9 genome editing strategy used to engineer the *ces-2(syb4992)* reporter expression. Bottom left, representative confocal micrographs of the expression pattern of *ces-2(syb4992[ces-2::SL2::GFP::H2B])* at L4 stage in the tail of both sexes. Bottom right, quantification of the number of CES-2::GFP-expressing cells from E, Bottom left. n=15 worms per group. Head pharynx and tail are outlined in white dashed line. Autofluorescence is indicated by yellow asterisk mark. Scale bars represents 10µm. Vertical bars in box-and-whiskers graph represent median with dots showing all points from min to max. We performed a Mann-Whitney test for each comparison, **** p < 0.0001.

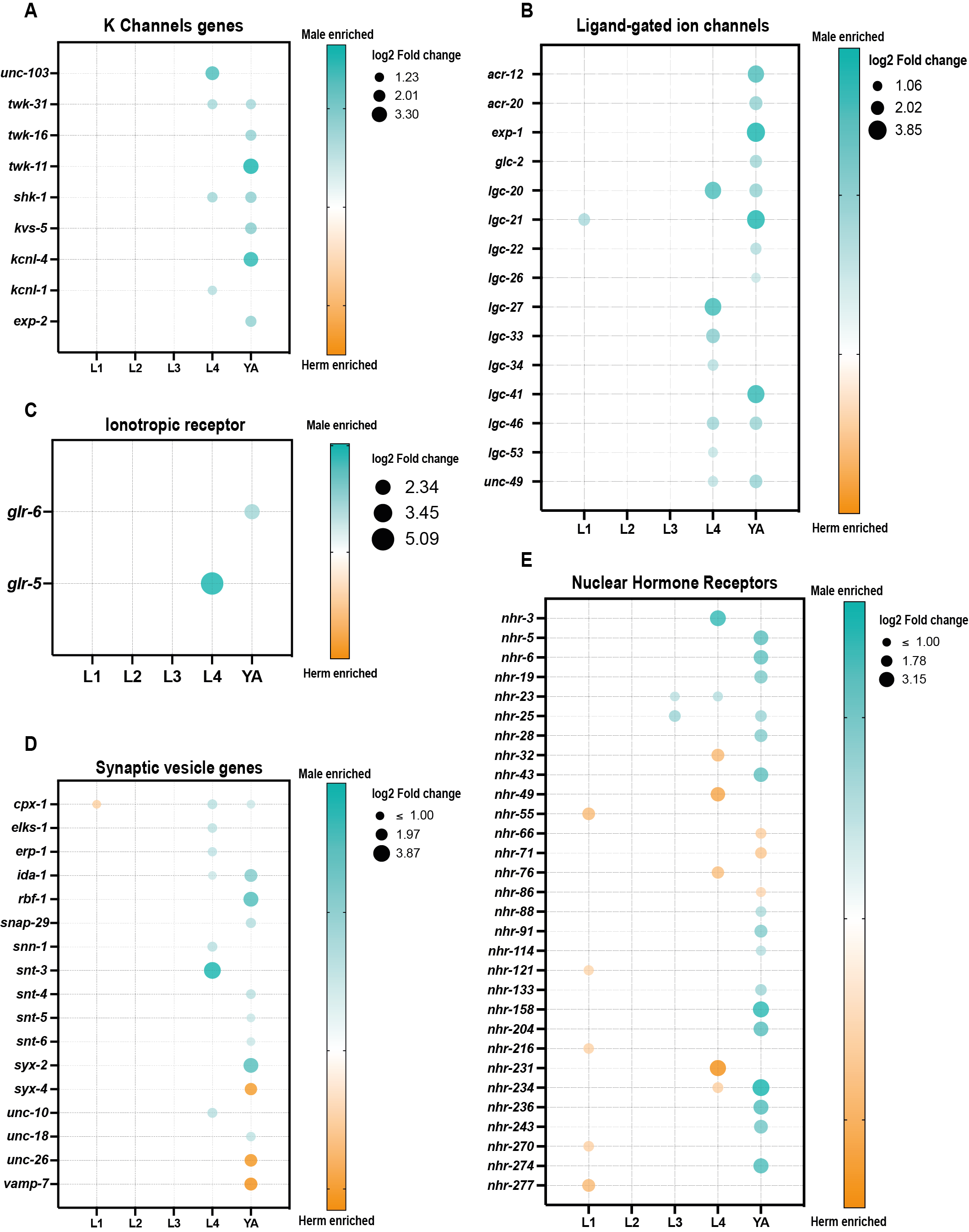

**Figure S5: Neuronal gene families with sexually dimorphic regulation**.

(A-E) Bubble plot representation of K channel (A), Ligand-gated ion channels (B), Ionotropic receptors (C), synaptic vesicle genes (D), and Nuclear hormone receptors (E) genes differentially expressed in any of the five developmental stages of the two sexes. Bubble size represents log2 of fold change in expression of that gene (only genes that passed the filter padj <= 0.05, |log2 fold change| >= 1 and basemean >= 5 are plotted), and bubble color represents enrichment in either sex (male enrichment in cyan, hermaphrodite enrichment in orange).

­­­­­
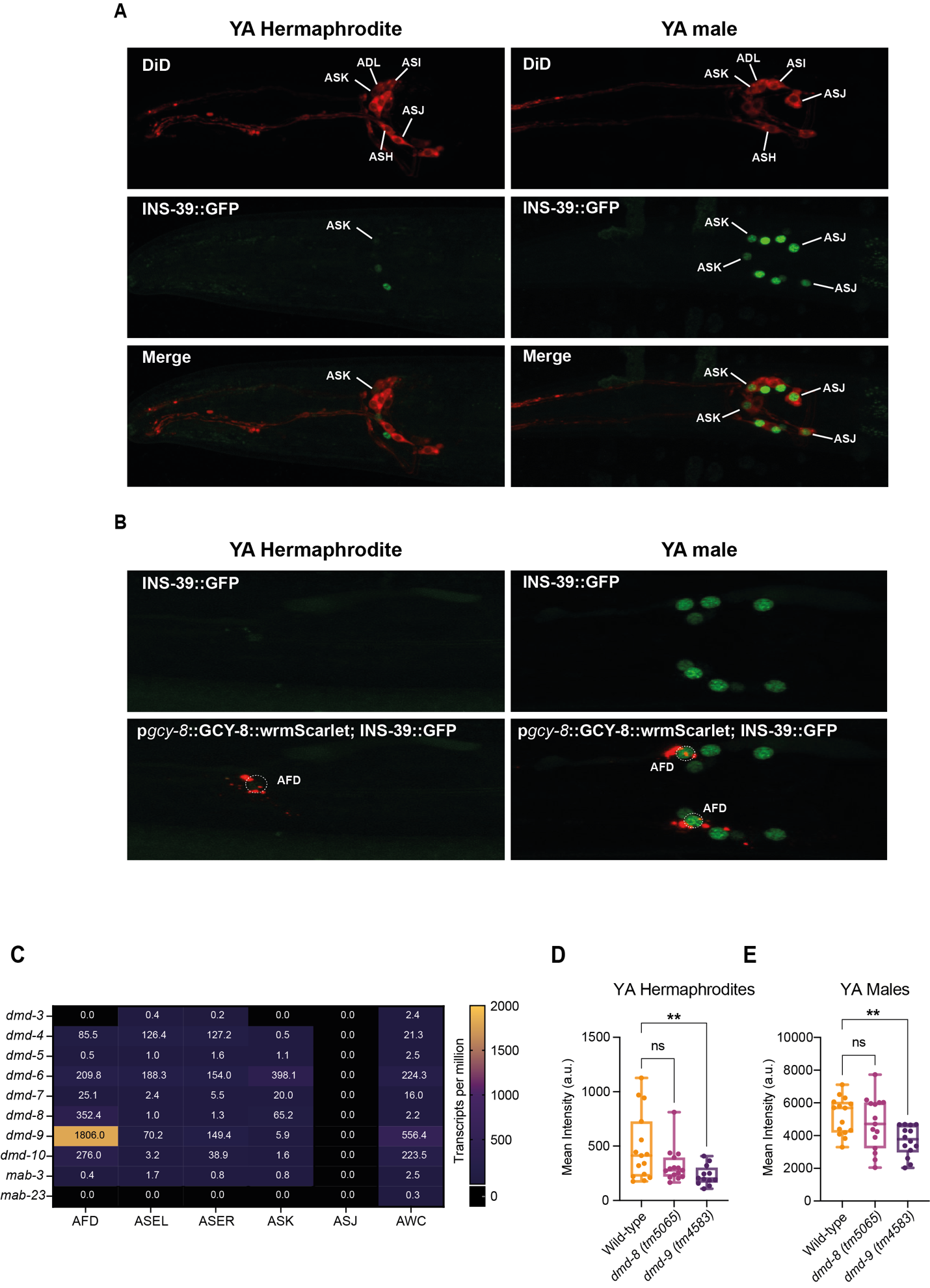
­­

**Figure S6: Identification of cells in which INS-39 is expressed.**

**(**A) Representative confocal micrographs of a young adult hermaphrodite (left) and male (right) expressing the INS-39::GFP reporter and co-stained with Vybrant lipophilic dye (DiD), enabling identification of sensory amphid neurons. Scale bar 10µm.

(B) Representative confocal micrographs of a young adult hermaphrodite (left) and male (right) expressing the INS-39::GFP reporter with AFD marker. Scale bar 10µm.

(C) Heatmap of gene expression pattern of all DMD genes in AFD, ASE, ASK, ASJ and AWC neurons from previously published single-cell/bulk-sorted neuronal transcriptomic data sets for DMD genes expression that coincide with *ins-39* expression pattern^2–4^.

(D-E) Quantification of mean GFP intensity in wild-type, *dmd-8* mutant (*tm5065*) and *dmd-9* mutant (*tm4583*) hermaphrodites (D) and male (E) using the *ins-39(syb4915)* reporter expression. n =12-15 worms per group. Vertical bars in box-and-whiskers graph represent median with dots showing all points from min to max. We performed Kruskal-Wallis test for (D) and (E) comparison, ** p < 0.01, ns-not significant.

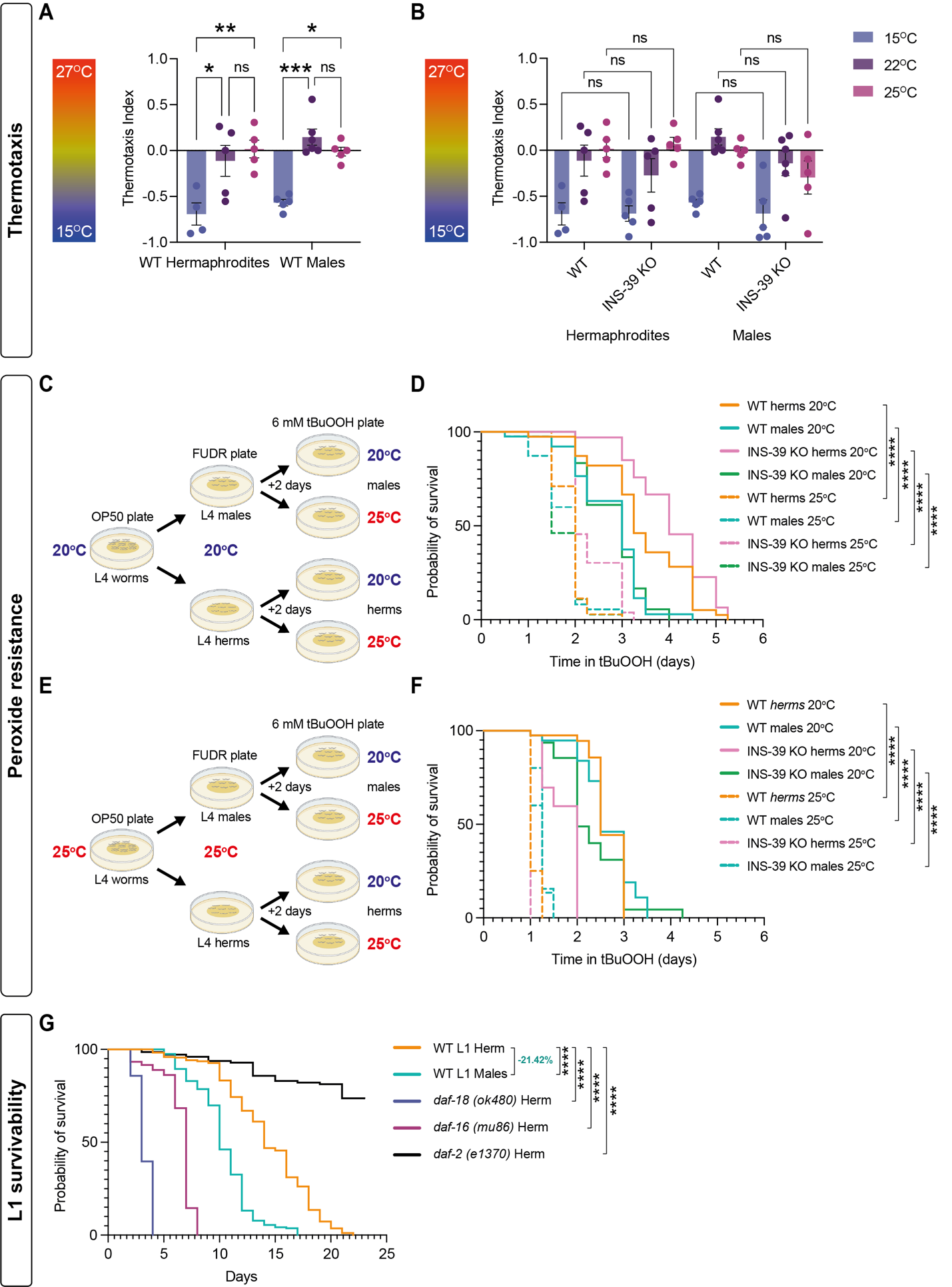

**Figure S7: Functional characterization of the role of INS-39.**

(A) Quantification of the thermotaxis index in wild-type hermaphrodites and males habituated at 15^o^C, 22^o^C and 25^o^C for 6 hours. Error bars are standard error of the mean (SEM). We performed a one-way ANOVA test for each comparison. Each dot represents independent biological repeats.

(B) Quantification of the thermotaxis index in wildtype and INS-39 KO CRISPR strain in hermaphrodites and males habituated at 15^o^C, 22^o^C and 25^o^C for 6 hours. Error bars are standard error of the mean (SEM). We performed a one-way ANOVA test for each comparison. Represent 4 biological replicates with n =50-120 worms per replicates.

(C) Schematic of the survival assay on 6mM tBuOOH for D.

(D) Cumulative survival graph on 6mM tBuOOH of wild-type and INS-39 KO hermaphrodites and males grown at 20°C and assayed 20°C or at 25°C. n =40 worms per group.

(E) Schematic of the survival assay on 6mM tBuOOH for E.

(F) Cumulative survival graph on 6mM tBuOOH of wild-type and INS-39 KO hermaphrodites and males grown at 25°C and assayed 20°C or at 25°C. n =40 worms per group.

(G) Cumulative L1 survival graph of wild-type hermaphrodites, wild-type males, *daf-18 (ok480)* hermaphrodites, *daf-16 (mu86)* hermaphrodites, *daf-2 (e1370)* hermaphrodites. See methods for n.

The statistical significance for survival was calculated using the Log-rank (Mantel-Cox) test. **** p < 0.0001, *** p < 0.001, ** p < 0.01, * p < 0.05, ns-non-significant.

**Description of Supplementary Tables**

**Supplementary Table S1**. Male percentages, raw read counts and RINe scores per sample.

**Supplementary Table S2**. Raw and normalized reads count of male and hermaphrodite genes per sample. Male-enriched and hermaphrodite-enriched genes with their human ortholog and their associated Online Mendelian Inheritance in Man (OMIM) human-disease phenotypes.

**Supplementary Table S3**. Hermaphrodite and male enriched genes with stage-specific or stage-shared regulation.

**Supplementary Table S4**. Male-enriched and hermaphrodite-enriched transcriptional factor, homeodomain, GPCR, neuropeptide, DM domain, K-channel, ligand-gated ion channels, ionotropic receptor, synaptic vesicle genes and nuclear hormone receptors.
